# High throughput screening to identify inhibitors of the type I interferon – major histocompatibility complex class I pathway in skeletal muscle

**DOI:** 10.1101/2020.03.20.998997

**Authors:** Travis B. Kinder, Patricia Dranchak, James Inglese

## Abstract

Immunosuppressants used to treat autoimmunity are often not curative and have many side effects. Our purpose was to identify therapeutics for autoimmunity of the skeletal muscle termed idiopathic inflammatory myopathies (myositis). Recent evidence shows the pro-inflammatory type I interferons (IFN) and a down-stream product major histocompatibility complex (MHC) class I are pathogenic in myositis. We conducted quantitative high throughput screening on >4,500 compound titrations through a series of cell-based assays to identify those that inhibit the type I IFN-MHC class I pathway in muscle precursor cells (myoblasts). The primary screen utilized CRISPR/Cas9 genome-engineered human myoblast containing a pro-luminescent reporter HiBit fused to the C-terminus of endogenous MHC class I. Active compounds were counter-screened for cytotoxicity and validated by MHC class I immunofluorescence, Western blot, and RT-qPCR. Actives included Janus kinase inhibitors and epigenetic/transcriptional modulators. Testing in animal models and clinical trials is warranted to translate these therapies to myositis patients.

## Introduction

Autoimmunity is the result of a break in immune self-tolerance, where the body’s adaptive immune system mistakenly attacks host tissues. Autoinflammation is chronic sterile inflammation due to environmental or genetic triggers and mediated by the innate immune system (Park et al., 2012). The idiopathic inflammatory myopathies, collectively referred to as myositis, is a group of systemic autoimmune diseases that has characteristics of autoinflammation, with the common feature of skeletal muscle inflammation and weakness often accompanied by autoantibodies and pathologies of the skin, lungs, or vasculature (Dalakas, 2015). Their causes are not fully understood but thought to be a combination of genetic predisposition and environmental triggers (Miller et al., 2018). Histological features vary between subtypes, but can include skeletal muscle infiltration by inflammatory cells, muscle fiber degeneration/regeneration, atrophy, fibrosis, necrosis, or fatty deposition. Current therapy consists of the glucocorticoid prednisone, immunosuppressive drugs like azathioprine or methotrexate, immune-modulating biologics such as intravenous immunoglobulin or rituximab, and physical exercise (Oddis and Aggarwal, 2018). Although these therapeutics can reduce disease burden in many patients, they have serious side effects and often do not completely eliminate muscle inflammation or restore muscle function (Lundberg et al., 2000). Therefore, there is a need for more targeted, less toxic therapies with which to treat myositis.

A potential novel therapeutic target in myositis is the pro-inflammatory cytokine family of type I interferons (IFN). A growing body of evidence shows an association between the abundance of type I IFN, along with a signature of IFN-stimulated genes (ISGs), and myositis pathology (Gallay et al., 2019). Type I IFNs are a family of glycosylated 20 kDa cytokines that interfere with viral replication in host cells, inhibit cellular proliferation, and have been implicated in a number of autoimmune conditions (González-Navajas et al., 2012). In humans, there exists 16 type I IFNs: 12 IFN-α subtypes, IFN-β, IFN-ε, IFN-κ, and IFN-ω. There is evidence that IFN-β can be produced by muscle precursor cells called myoblasts (Tournadre et al., 2012), and the level of IFN-β protein, but not IFN-α or −ω, is correlated to ISGs in dermatomyositis (Liao et al., 2011). Promisingly, small pilot trials and anecdotal reports show that blocking type I IFN signaling with Janus kinase (JAK) inhibitors may lead to improvements in myositis patients (Ladislau et al., 2018; Paik et al., 2018; Papadopoulou et al., 2019).

Type I IFNs are potent inducers of major histocompatibility complex (MHC) class I expression, including in myoblasts (Tournadre et al., 2012). MHC class I bridges the innate and adaptive immune systems by presenting peptides to CD8+ T lymphocytes to coordinate cytotoxic T cell killing. Several genome-wide association studies have confirmed that the ancestral 8.1 haplotype of the human leukocyte antigen (HLA) region of the genome, encoding a number of immune genes including MHC classes I and II, confers the highest susceptibility to myositis (Miller et al., 2018). MHC class I is not expressed in healthy muscle fibers, but its expression is an early and prominent feature of myositis muscle (Li et al., 2004). Patient muscles exhibiting refractory weakness without inflammatory cell infiltration have been observed to have chronic MHC class I over-expression, even after prednisone treatment (Lundberg et al., 2000; Nyberg et al., 2000). Additionally, transgenic over-expression of MHC class I in mouse skeletal muscle recapitulates many features of myositis, including endoplasmic reticulum (ER) stress and up-regulation of ISGs (Kinder et al., 2020; Nagaraju et al., 2000). Taken together, chronic expression of MHC class I, or certain alleles of it, may be pathogenic in skeletal muscle and could be a therapeutic target in myositis (**Figure 1A**).

**Figure 1.**
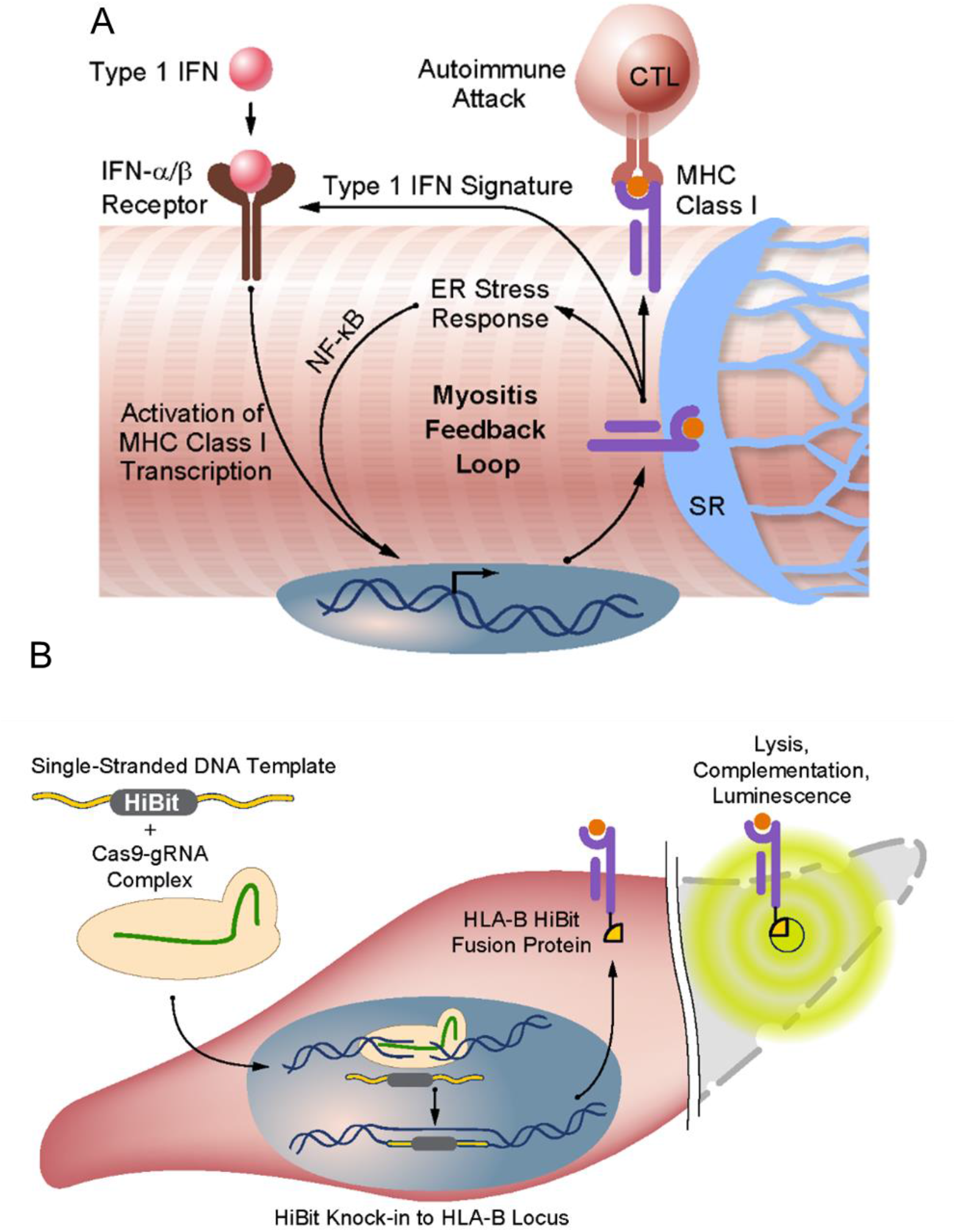
Illustrations of the type 1 IFN – MHC class I pathway in skeletal muscle and the creation of an MHC class I (HLA-B)-HiBit luminescent reporter fusion protein using Cas9 genome-editing in myoblasts. **A,** Type 1 IFNs activate the interferon-α/β receptor on the cell surface, which drives the expression of IFN responsive genes including MHC class I. In skeletal muscle, MHC class I expression leads to an ER stress response, further activation of the IFN signature, and autoimmune attack by cytotoxic T lymphocytes (CTLs). **B,** For high throughput screening assays, the HiBit reporter peptide was fused to the C-terminus of endogenous HLA-B in immortalized human myoblasts by transfecting single-stranded DNA template for HiBit, flanked by homology arms, and Cas9 protein-guide RNA (gRNA) complex. HLA-B expression could then be measured using a lytic luminescence assay via complementation of HiBit with the LgBit subunit from NanoLuc plus furimazine substrate.

In order to find new therapeutics that could reduce autoimmunity and inflammation in myositis, we have developed a series of *in vitro* assays to identify small molecule compounds that inhibit the type I IFN – MHC class I pathway in skeletal muscle. We used CRISPR/Cas9 genome-engineering of human myoblasts to fuse a small, pro-luminescent Nano luciferase sequence (called HiBit) to the C-terminus of endogenous MHC class I (**Figure 1B**) and performed quantitative high throughput screening (qHTS) to test >4,500 compounds with concentrations spanning several orders of magnitude. Active compounds were followed-up by HLA-ABC immunofluorescence (IF), reverse transcriptase quantitative PCR (RT-qPCR), and Western blot. Several broad classes of active compounds were identified including JAK inhibitors, assorted non-JAK kinase inhibitors, and epigenetic/transcriptional modulators, which require further testing in animal models and clinical trials before translation to myositis patients.

## Materials and Methods

### Cell culture

Immortalized human myoblast cell line C25cl48 was obtained from Vincent Mouly, PhD (Institut de Myologie, Paris, France), which was derived from the semitendinosus muscle of a healthy 25 year-old male and transduced with both telomerase and cyclin-dependent kinase 4 retroviruses (Mamchaoui et al., 2011). All reagents were from Thermo Fisher, Waltham, MA, unless otherwise noted. C25cl48 growth medium (GM) comprised 4:1 ratio DMEM Glutamax (10566) : M199 (11150) + 10% FBS (Hyclone Standard, GE, Chicago, IL, SH30088) + 1% Pen/Strep (15140122) + 50 ug/mL fetuin + 10 ng/mL hEGF + 1 ng/mL bhFGF + 10 ug/mL insulin + 0.4 ug/mL dexamethasone (Promocell, Heidelberg, Germany, C-39360), and cells were maintained at 37 °C, 5% CO_2_, and 95% relative humidity. Cells were periodically assayed for mycoplasma contamination and were negative throughout these experiments. For all assays, cells were plated without dexamethasone. Recombinant human IFN-β (Peprotech, Rocky Hill, NJ, 300-02BC) was used to stimulate MHC class I expression.

### CRISPR/Cas9 genome-engineering

The strategy for genome-engineering to tag HLA-B with the HiBit peptide was developed based on a previous publication from Promega Corporation, Madison, WI (Schwinn et al., 2018). Karyotyping of C25cl48 cells was performed by the Cytogenetics and Microscopy Core, NHGRI, NIH (Bethesda, MD) to confirm normal genome structure, and high resolution HLA typing was performed by Johns Hopkins Immunogenetics Lab (Baltimore, MD) to determine HLA alleles for editing (**Supplementary Figure 1**). CRISPR RNA (crRNA) (GTGTCTCTCACAGCTTGAAAAGG) was designed and purchased using Integrated DNA Technologies (IDT, Coraliville, IA) Custom Alt-R CRISPR-Cas9 guide RNA online tool (www.idtdna.com) to target 2 bp from the terminal stop codon of HLA-B*08:01:01:01. crRNA was complexed with Alt-R CRISPR-Cas9 tracrRNA (IDT, 1072533) to form the guide RNA (gRNA). Donor single-stranded DNA (ssDNA) containing a 33 bp HiBit sequence (GTGAGCGGCTGGCGGCTGTTCAAGAAGATTAGC) and flanked by 70 bp homology arms was synthesized as an Ultramer Oligo with standard desalting (IDT). 1.3×10^5 C25cl48 cells/reaction were electroporated with a ribonucleoprotein consisting of 125 pmol TrueCut Cas9 Protein v2 (A36498) + 150 pmol gRNA + 120 pmol ssDNA using P5 Primary Cell 4D-Nucleofector X Kit S (Lonza, Walkersville, MD, V4XP-5032) in a 16-well Nucleocuvette Strip and the Amaxa 4D-Nucleofector X Unit (Lonza) running program EY-100. Clones were isolated by limiting dilution in a 96-well plate (0.6 cells/well).

### HLA-B HiBit and CellTiter-Glo viability assays

300 HLA-B HiBit cells/well were plated into white 1536-well plates (Greiner, Monroe, NC, 789173-F) in 5 uL/well of GM using a Multidrop Combi (Thermo Scientific). Column 1 did not contain IFN-β as a control, and all other columns contained 4.0 ng/mL IFN-β, where column 4 was used as IFN-β only control. 23 nL/well DMSO was added to columns 1-4 while 23 nL/well compounds in DMSO were added to columns 5-48 using a pintool (Wako, San Diego, CA). For counter screening, digitonin (Sigma, St Louis, MO, D141) in DMSO was added to column 2 at a final concentration of 92 uM as a cytotoxic control. For Nano-Glo HiBit Lytic Detection System (HiBit assay) (Promega, N3040) and CellTiter-Glo Luminescent Cell Viability Assay (CTG) (Promega, G7572), 2.5 uL/well reagent was added with a BioRAPTR FRD (Beckman Coulter, Sykesville, MD), plates were incubated in the dark at ambient temperature for 10 min, and luminescence measured with a ViewLux 1430 Ultra HTS (Perkin Elmer, Waltham, MA) running Wallac 1430 Manager and Explorer software v3.02. Two of the three primary screening libraries (NPC and MIPE5.0) were pinned, incubated, and assayed on our in-house Kalypsys robotics system. IFN-β titration in Figure 2 was dispensed at 50 nl/well with a Mosquito (TTP Labtech, Boston, MA). Refer to supplemental materials and methods for detailed protocol tables as we have described previously (Inglese et al., 2007b).

**Figure 2.**
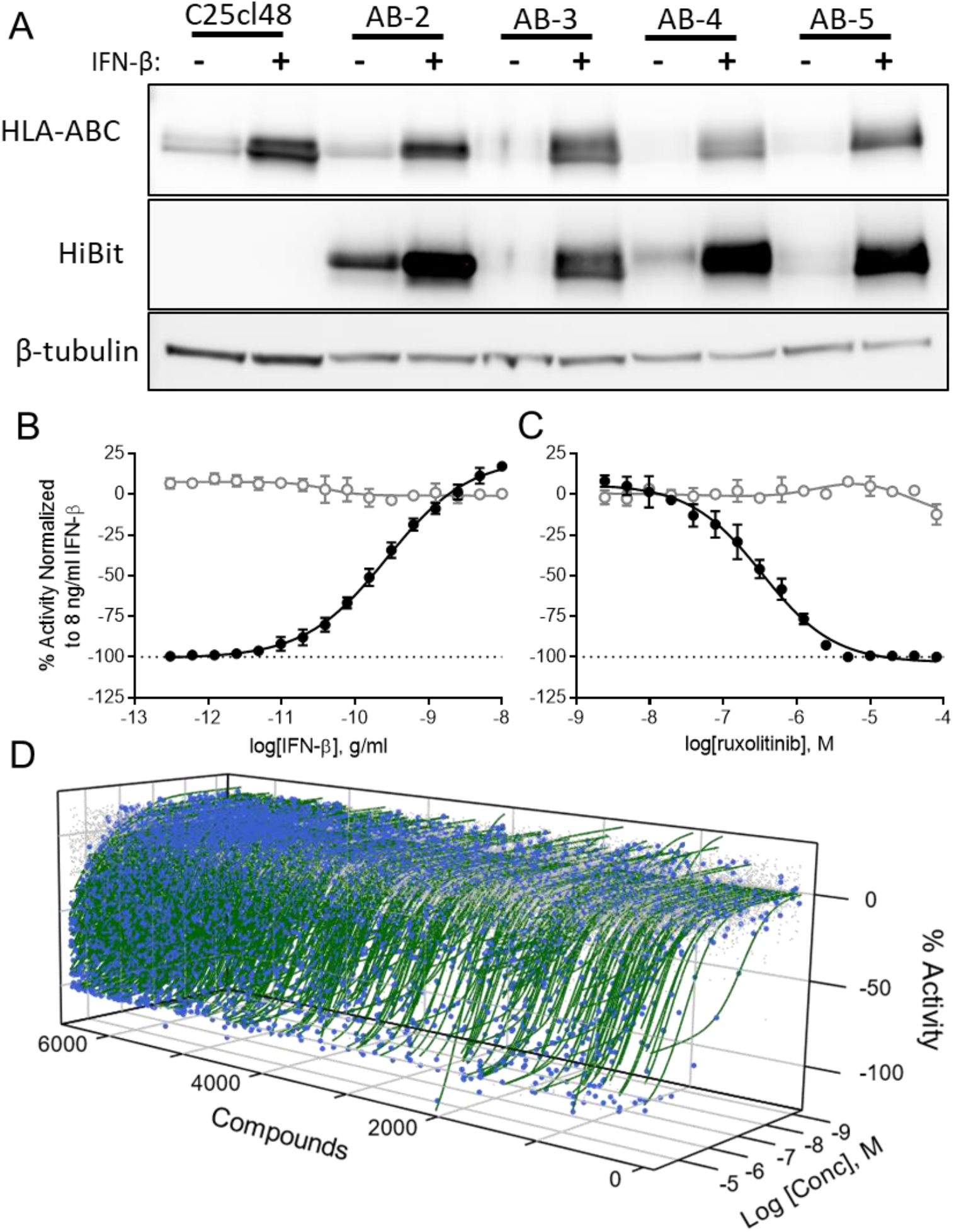
Assessment of HLA-B HiBit genome-engineered myoblast clones and primary screening results. **A,** Western blot analysis of MHC class I (HLA-ABC), HiBit, and β-tubulin loading control in parental C25cl48 cells and several genome-engineered clones treated ± 8.0 ng/mL IFN-β for 24 hrs showed up-regulation of endogenous HLA-ABC in all cells and HiBit up-regulation in genome-engineered clones AB-2, -3, -4, and -5. **B** and **C,** HiBit assay luminescence CRCs for control treatments in 1536-well plates. Mean ± standard deviation of triplicate experiments. Blacked filled circles – HiBit, open grey circles – CTG. **B,** 16-point, 1:2 titration of IFN-β starting at 10 ng/ml for 24 hrs. AC50 = 270 ± 22 pg/ml **C,** Cells treated with 4.0 ng/ml IFN-β plus a 16-point, 1:2 titration of JAK inhibitor ruxolitinib starting at 80 uM for 24 hrs. IC50 = 340 ± 36 nM. **D,** Waterfall plot showing all data from primary screening of 6,576 compounds at 7 to 11 titrations each. Gray dots are data from inactive compounds, blue dots with green curves are data from active compounds selected for follow-up. Data was normalized within each plate to 4.0 ng/ml IFN-β = 0% and DMSO alone = −100% activity.

### HLA-ABC IF secondary assay

Assays were performed in black/clear bottom 1536-well plates (Aurora, Scottsdale, AZ, 00019324). 25 nL/well DMSO was dispensed into columns 1, 2, & 4, 20 mM digitonin in DMSO was dispensed into column 3, and compounds in DMSO were dispensed into columns 5-48 via acoustic dispensing (Labcyte Echo 655, San Jose, CA). Then, 125 C25cl48 cells/well were added in 5 uL/well GM using a Multidrop Combi. Columns 1 & 2 did not contain IFN-β as a control, and all other columns contained 4.0 ng/mL IFN-β, where column 4 was used as IFN-β only control. See Supplementary Materials and Methods for IF protocol. Plates were analyzed by laser scanning cytometry using an Acumen eX3 (TTP Labtech) running Cellista v4.3.4.0 software and utilizing 405, 488, and 561 nm lasers. HLA-ABC+ cells were quantified as number of composite objects positive for Hoechst & HLA-ABC & CellMask Red / cell count. Cell count was quantified as number of objects positive for CellMask Red. For each compound plate, triplicate plates were assayed and averages were plotted as concentration-response-curves (CRCs). Representative microscopy images of control wells were taken on an IN Cell Analyzer 2200 instrument and software v7.1 (GE).

### RT-qPCR assays

Assays were performed in 384-well plates (Corning, NY, 3765). 150 nL/well DMSO was dispensed for 6 wells per control, and compounds in DMSO were dispensed via acoustic dispensing (Labcyte Echo 655). Then, 780 C25cl48 cells/well were added in 30 uL/well of GM using a multichannel pipette. 6 wells did not contain IFN-β as a control, and all other wells contained 4.0 ng/mL IFN-β, where 6 wells were used as IFN-β only control. Cells were harvested and assayed using the Cells-to-C_T_ 1-Step TaqMan Kit (A25602) and TaqMan assays for *HLA-A* (Hs01058806_g1), *HLA-B* (Hs00741005_g1), *HLA-C* (Hs00740298_g1), *IFIT1* (Hs03027069_s1), *ISG15* (Hs01921425_s1), *MX1* (Hs00895608_m1), and *HPRT1* (Hs02800695_m1). All assays utilized FAM reporter, except HPRT1 utilized VIC reporter for multiplexing. After incubation with compound, cells were washed with PBS and lysed in 13 uL lysis buffer. 1.4 uL lysate/reaction was PCR amplified in 14 uL master mix using a ViiA7 instrument (Life Technologies) running ViiA7 software v1.0, and expression levels were calculated using the ΔΔC_T_ method relative to *HPRT1*.

### Western blotting

200,000 cells/well were plated into 6-well plates in 2 mL/well GM, treated the next day at the IC95 according to HiBit CRCs, washed with PBS, and lysed in 75 uL/well RIPA buffer + protease inhibitor (Sigma, R0276 and P8340) + PhoSTOP (Roche, Basel, Switzerland, 04906845001). See Supplementary Materials and Methods for detailed protocols.

### Whole Genome Sequencing (WGS)

Conducted by the Center for Applied Genomics at the Children’s Hospital of Philadelphia, PA. See Supplementary Materials and Methods for detailed protocol and data analysis methods.

## Results

### Development of HLA-B HiBit reporter cells for qHTS

Nine genome-engineered HiBit clones were treated with IFN-β and assayed by Western blot to estimate the molecular weight and expression level of MHC class I and HiBit (**Figure 2A** and **Supplementary Figure 2**). IFN-β treatment up-regulated endogenous MHC class I (HLA-ABC) protein in parental C25cl48 cells and edited clones, and up-regulated HiBit in all edited clones. Additionally, the molecular weight for the HiBit band (∼45 kDa) was identical to that for HLA-ABC, providing evidence of proper on-target integration. HiBit signal did not appear in any other location, indicating no off-target integration in the expressed genome (**Supplementary Figure 3**).

Clone AB-7 was chosen for qHTS (referred to as HLA-B HiBit cells) due to its superior signal-to-background ratio of 32.5 and Z’-factor of 0.89 (**Supplementary Table 1**), a parameter to assess HTS assays’ dynamic range that takes into account the difference between positive and negative controls and the variability of each (Zhang et al., 1999). Pharmacological characterization of this clone demonstrated a sigmoidal response to IFN-β yielding an AC50 of 270 ± 22 pg/mL (**Figure 2B**). Reported bioactivity of IFN-β is approximately 100 pg/ml in a TF-1 proliferation assay (Mire-Sluis et al., 1996). The CTG viability assay showed a concentration-response inhibition of cell proliferation with increasing concentration of IFN-β, as reported in other cell types (González-Navajas et al., 2012). HLA-B HiBit cells stimulated with 4.0 ng/mL IFN-β and titrated with the JAK inhibitor ruxolitinib yielded a sigmoidal response with an IC50 of 340 ± 36 nM (**Figure 2C**). These results are in agreement with literature values of ruxolitinib inhibition for IL-6 stimulation of whole blood with an IC50 = 282 ± 54 nM (Quintás-Cardama et al., 2010). The CTG viability assay for IFN-β + ruxolitinib titration showed a bell-shaped curve with increased cell viability peaking at 5 uM and then decreasing below baseline at 80 uM. Increased viability was observed at effective concentrations of drug, which is likely due to blocking the anti-proliferative effects of IFN-β, thus restoring cell growth. Then at higher concentrations, it becomes toxic and decreases viability.

To thoroughly investigate on- and off-target insertion of HiBit, WGS was performed on C25cl48 cells and four genome-engineered clones (AB-3, -5, -7, and -11). The average depth of sequencing coverage was 39.4 reads (**Supplementary Table 2**), and the resulting sequences are publicly available on the Sequence Read Archive (accession: PRJNA608961). Sequence analysis verified HiBit integrated only once at the C-terminus of *HLA-B* and nowhere else in the genome (**Supplementary Table 3**). Phase analysis showed HiBit fused only to the targeted 08:01 allele and not the 18:01 allele (**Supplementary Figure 4**), meaning a heterozygous integration. Integration into clones AB-3 and -5 had exact sequence matches for HiBit, but for clones AB-7 and -11, small genomic portions of sequence surrounding HiBit were duplicated, representing a duplication of exon 7 coding for 14 amino acids of the cytoplasmic tail. These sequencing results were obtained after primary screening with AB-7, and follow-up assays validated those results, so the duplication did not appear to alter the expression of HLA-B.

### HLA-B HiBit primary screening

Three libraries were screened containing a total of 6,576 compounds at 7 to 11 titrations each spanning the nM to uM range, and due to partial sample redundancy resulted in 4,679 unique compounds tested (**Supplementary Table 4**). CRCs were fit to all data and compounds were categorized based on efficacy and potency as previously described (Inglese et al., 2006). Briefly, curve class 1 is a complete curve with both asymptotes, 2 is an incomplete curve with one asymptote, 3 has a single point activity, and 4 is inactive. Additionally, an ‘a’ or ‘b’ suffix represents high efficacy of ≥ 80% or lower efficacy of < 80%, respectively, and negative curves are inhibitory while positive are stimulatory. Most compounds tested were inactive (4,945, 75% of total). We selected active compounds with curve classes -1a, -1b, and -2a (650, 14% of unique compounds) for follow-up (**Figure 2D**).

### HLA-B HiBit retest and CTG counter screen

Upon retesting by HLA-B HiBit assay, most samples reconfirmed inhibitory activity (568, 87%). Cellular viability was estimated by CTG, and about half the compounds (353, 54%) displayed some degree of cytotoxicity. To eliminate cytotoxic compounds, we compared the HiBit and CTG curves for each. Compounds were selected to move forward that had active HiBit curve classes of -1 and -2 with CTG IC50 >4-fold higher than the HiBit IC50, which resulted in 216 compounds. The CRCs for twelve of these compounds, representative of the different active classes, are shown in **Figure 3**.

**Figure 3.**
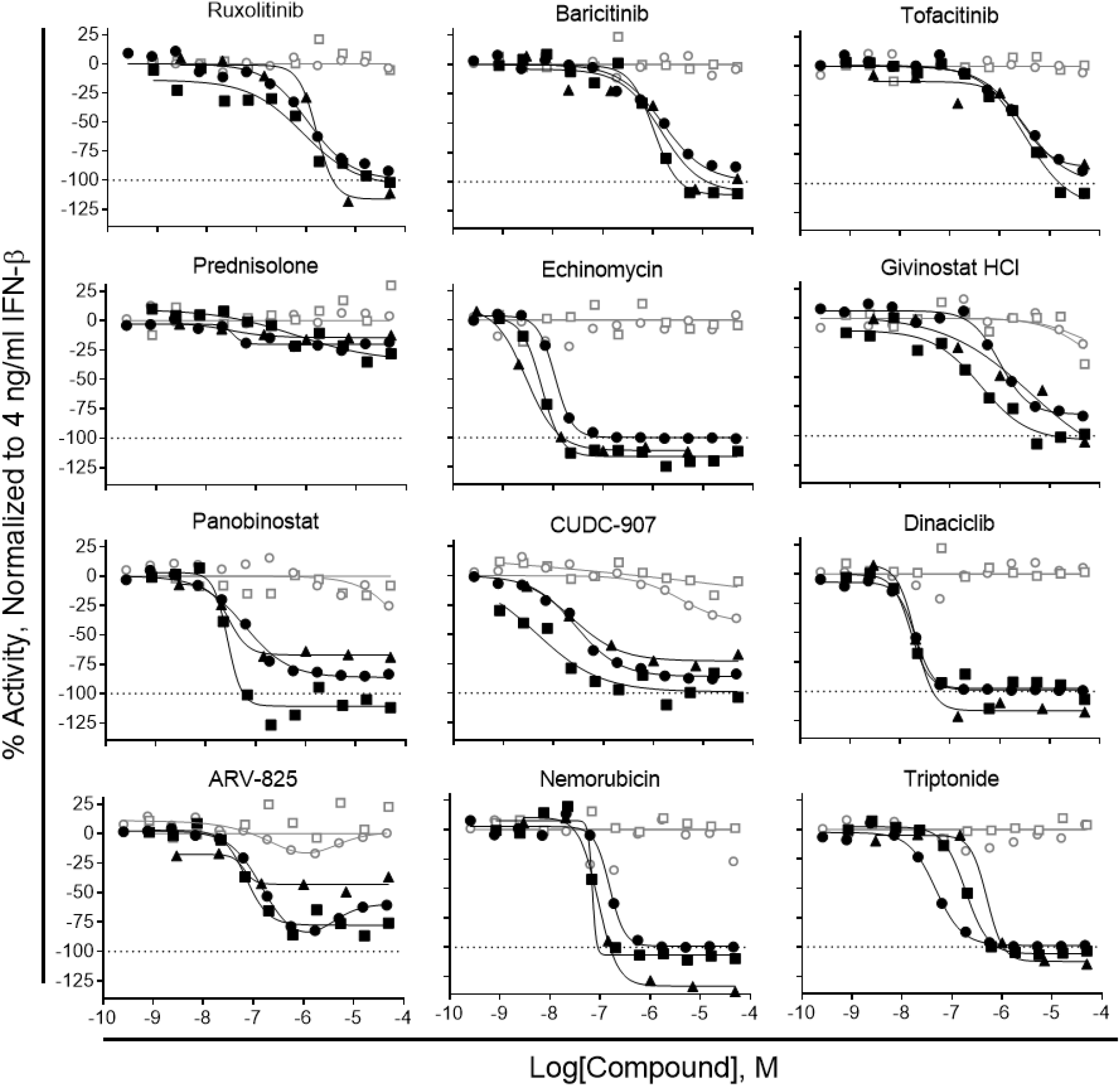
CRCs for representative active compounds. HLA-B HiBit clone AB-7 cells plated in 1536 well plates and treated for 24 hours with 4.0 ng/ml IFN-β plus titrations of active compounds from primary screening libraries. Black circles – HiBit, open gray circles – CTG, black boxes – IF HLA-ABC+, open gray boxes – IF cell count, black triangles – *HLA-B* RT-qPCR. For HiBit, IF, and RT-qPCR efficacy assays, data was normalized within each plate to 4.0 ng/ml IFN-β = 0% and DMSO alone = −100% activity. For CTG and cell count cytotoxicity assays, data was normalized within each plate to 4.0 ng/ml IFN-β = 0% and 92 uM Digitonin = −100% activity.

### HLA-ABC IF secondary assay

To confirm the actives biologically inhibited MHC class I protein production, rather than artifactually interfering with reporter signal (Thorne et al., 2010) or results being skewed by genome-engineering, we assayed follow-up compounds by HLA-ABC IF in parental C25cl48 cells. Fluorescence staining was quantified using laser scanning cytometry as we’ve described previously (Auld et al., 2006). Representative microscopy and cytometry images are shown in **Figure 4A**. The quantification of cells positive for HLA-ABC over titrations of IFN-β produced a CRC with an AC50 of 58 ± 7 pg/ml and had a growth inhibition effect as seen previously (**Figure 4B**). In the same HLA-ABC characterization experiments, IFN-β + ruxolitinib titration had an IC50 of 3,500 ± 630 nM with a bell shaped viability curve, where cells were proliferating in the presence of effective concentrations of ruxolitinib and then started to die at higher concentrations approaching 80 uM (**Figure 4C**). By this assay, 179 compounds (83%) displayed inhibitory activity. Most active compounds could now be grouped into several broad classes of inhibitors to: kinases, histone deacetylase (HDAC), DNA topoisomerase II, transcription factors such as nuclear factor κ-light-chain-enhancer of activated B cells (NF-κB), glucocorticoids, bromodomain-containing protein 4 (BRD4), proteasome, heat shock protein 90 (HSP90), and Na+/K+ ATPase. For follow-up, several representatives from each target group that had active HLA-ABC curve classes of -1 with cell count IC50 >3-fold higher than HLA-ABC IC50 were selected, resulting in 62 compounds. The HLA-ABC IF derived CRCs for the twelve representative compounds shown previously are displayed in **Figure 3**.

**Figure 4.**
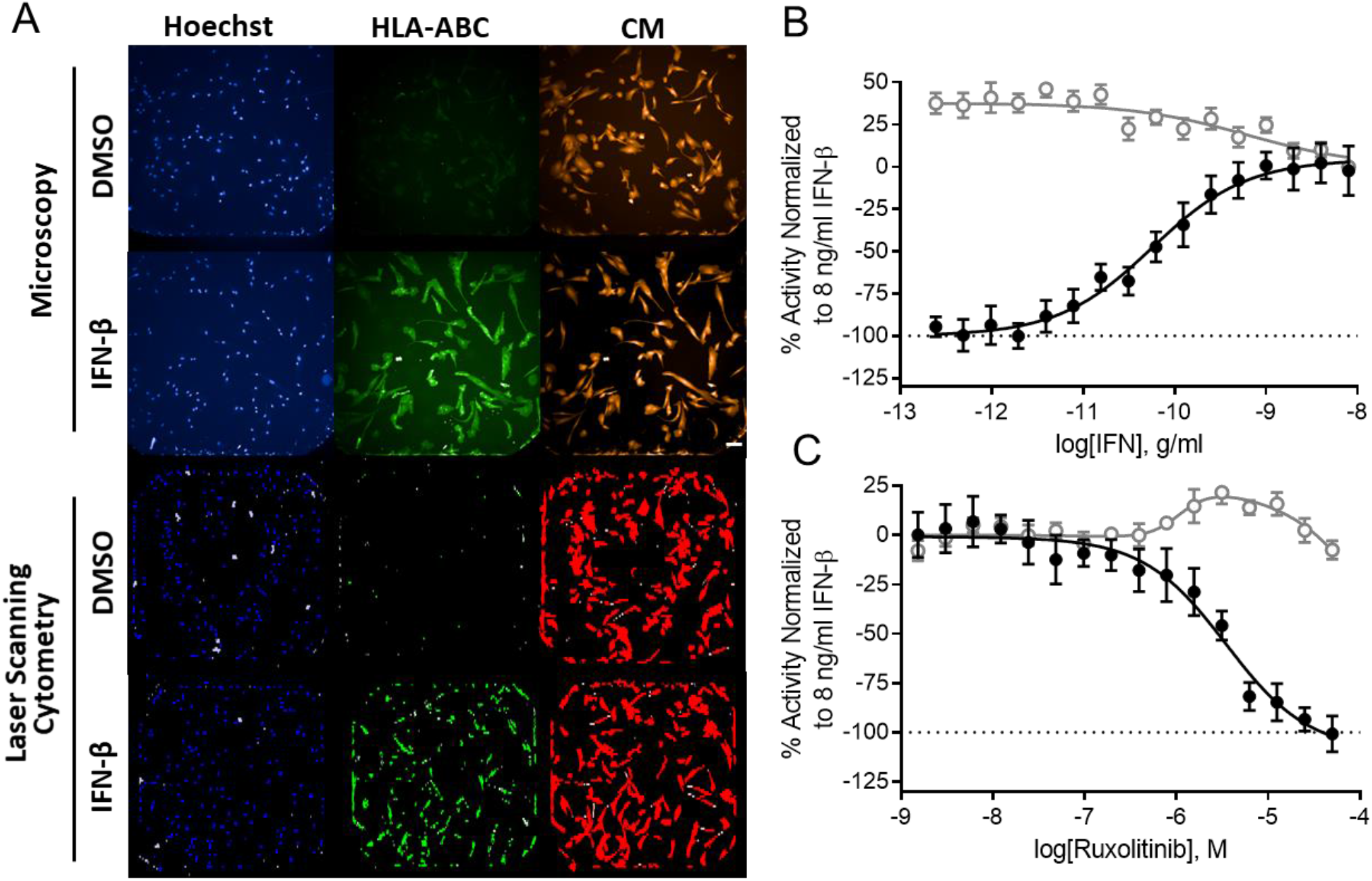
HLA-ABC IF analysis. **A,** Images of control wells for HLA-ABC IF in 1536 well plates. On top are fluorescent microscopy images at 10x magnification; white scale bar = 100 um. On bottom are laser scanning cytometry images. Treatment with IFN-β increased HLA-ABC protein expression in C25cl48 cells and can be detected by both technologies. **B** and **C,** Quantification of HLA-ABC+ cells for control treatments in 1536-well plates. Mean ± standard deviation of nine plates. Black filled circles – HLA-ABC+ cells, grey open circles – cell count. **B,** 16-point, 1:2 titration of IFN-β starting at 8 ng/ml for 48 hrs. AC50 = 58 ± 7 pg/ml. **C,** Cells treated with 8.0 ng/ml IFN-β plus a 16-point, 1:2 titration of JAK inhibitor ruxolitinib starting at 50 uM for 48 hrs. IC50 = 3,500 ± 630 nM. For HLA-ABC+ cells, data was normalized within each plate to 8.0 ng/ml IFN-β = 0% and DMSO alone = −100% activity. For cell count, data was normalized within each plate to 8.0 ng/ml IFN-β = 0% and 92 uM Digitonin = −100% activity.

### HLA-B RT-qPCR tertiary assay

To distinguish compounds that inhibit the pathway at the stage of *HLA-B* mRNA transcription versus those that decrease protein abundance, actives were assayed by RT-qPCR. By this assay, 57 compounds retained inhibitory activity (92%). The CRCs for the twelve representative active compounds are displayed in **Figure 3**. Notably, five targets that were active in HiBit and IF assays but were inactive or far less active by RT-qPCR were BRD4, proteasome, HSP90, Na+/K+ ATPase, and SERCA.

### Effect of active compounds on type I IFN pathway

To investigate mechanisms of action, Western blots were performed to determine the phosphorylation status of transcription factor STAT1 and RT-qPCR was performed for *HLA-A*, *-C* and three ISGs: *IFIT1*, *ISG15*, and *MX1*. Stimulation of the IFN-α/β receptor (IFNAR) results in JAK phosphorylation of STAT1/2, which then translocate to the nucleus and activate transcription of HLA genes and ISGs. For C25cl48 cells In the presence of IFN-β, STAT1 was phosphorylated between 0.5-16 hrs, whereas total STAT1 and HLA-ABC were up-regulated at 6.0-24 hrs (**Supplementary Figure 5**). Of the 11 representative actives tested by Western blot, only the JAK inhibitors ruxolitinib, baricitinib, and tofacitinib prevented phosphorylation of STAT1, but all compounds inhibited HLA-ABC protein expression with similar efficacies seen in previous experiments (**Figure 5A**). By RT-qPCR, IFN-β up-regulated *HLA-A* (3.1-fold), *HLA-C* (11-fold), *IFIT1* (100-fold), *ISG15* (100-fold), and *MX1* (3100-fold) (**Figure 5B**). These control samples assayed during screening showed IFN-β up-regulated *HLA-B* (42-fold). Compared to IFN-β treated cells, the JAK inhibitors decreased all genes tested >99% except *HLA-A*, which was only minimally up-regulated by IFN-β alone. For the HDAC inhibitors, givinostat normalized *HLA-A* and *-B* to DMSO levels while decreasing ISGs ∼90%, and panobinostat decreased all genes ∼80% with no effect on *HLA-A*. Echinomycin, nemorubicin, and triptonide normalized all genes. The dual phosphoinositide 3-kinases (PI3K) / HDAC inhibitor CUDC-907 decreased *HLA-C*, *IFIT1*, and *MX1 ∼*75%. BRD4 inhibitor ARV-825, which was efficacious in HiBit and IF assays, but had some cytotoxicity and was only marginally effective at reducing *HLA-B* mRNA decreased *IFIT1* 84%, but increased *ISG15* 2.8-fold. Cyclin-dependent kinase 2/4 (CDK2/4) inhibitor dinaciclib increased *HLA-C*, *ISG15,* and *MX1 ∼*3.3-fold. Prednisolone increased *ISG15* and *MX1 ∼*3.4-fold. Taken together, these data indicate that JAK inhibitor drugs block the type I IFN – MHC pathway at the stage of STAT phosphorylation, while the other active compounds block the pathway through non JAK-STAT mechanisms. 11/12 compounds effects on ISGs were comparable to their effects on *HLA-B* mRNA measured in qHTS, where the JAK inhibitors, echinomycin, nemorubicin, and triptonide had >99% efficacy in reducing all HLA and ISG genes. Givinostat, panobinostat, and CUDC-907 had about 70-80% efficacy for reducing all HLA and ISG genes. ARV-825 had low efficacy in reducing mRNA for any gene. The exception to these trends was the CDK2/4 inhibitor dinaciclib, which was highly effective at reducing *HLA-B* mRNA but did not decrease any of the other genes tested, which warrants further investigation into this selective mechanism of action.

**Figure 5.**
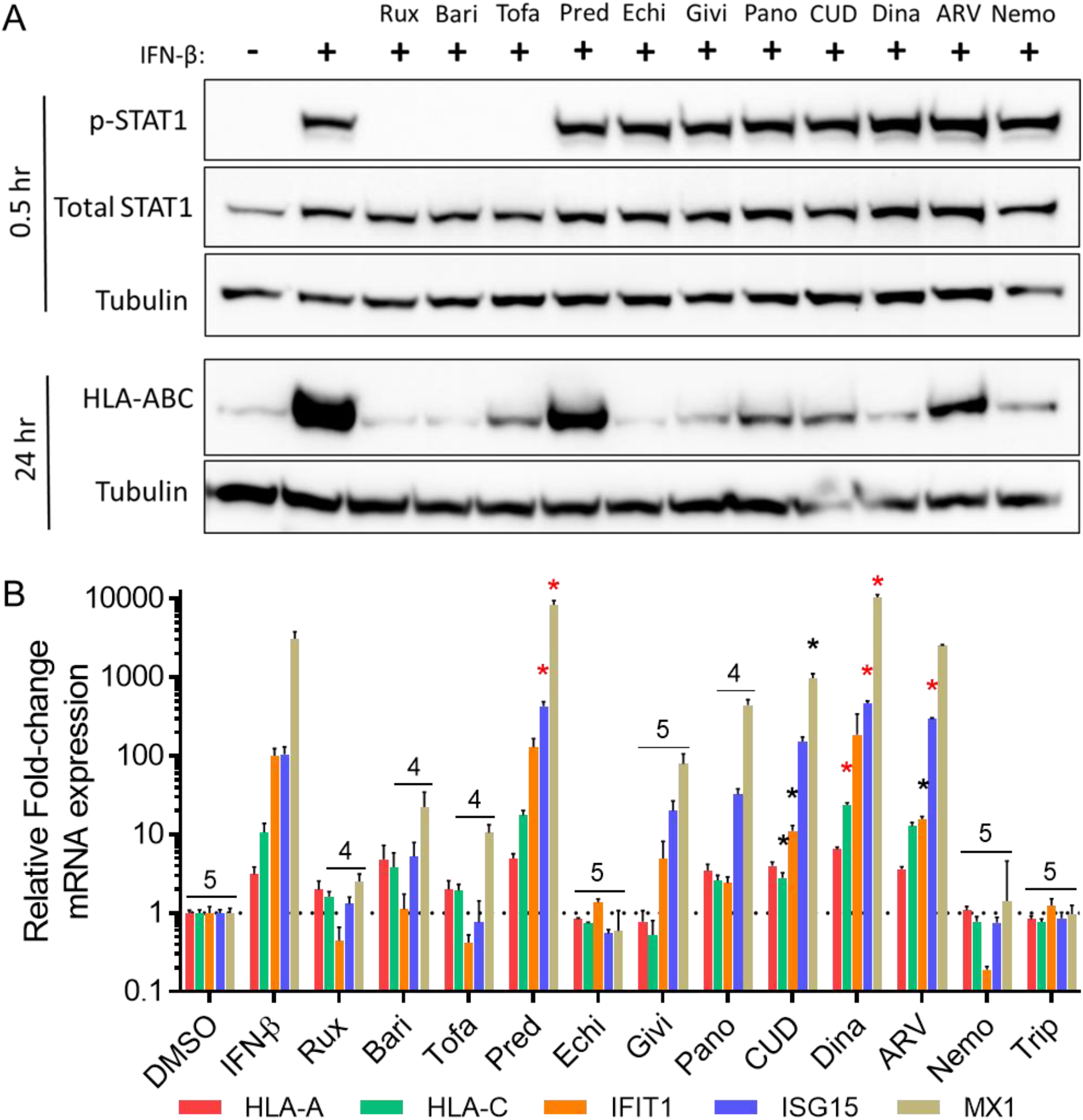
Effects of representative active compounds on Type I IFN pathway in myoblasts. **A,** Western blot analysis of C25cl48 cells’ STAT1 activation by phosphorylation after 0.5 hrs of treatment and HLA-ABC expression after 24 hrs of treatment with 4.0 ng/ml IFN-β plus the IC95 concentration of active compounds. Only the three JAK inhibitors ruxolitinib (rux), baricitinib (bari), and tofacitinib (tofa) prevented phosphorylation of STAT1, whereas all active compounds reduced HLA-ABC expression with similar efficacies seen in previous experiments. **B,** RT-qPCR analysis of *HLA-A, HLA-B, IFIT1, ISG15*, and *MX1* relative to *HPRT1* in C25cl48 cells treated with 4.0 ng/ml IFN-β plus 50 uM compound for 24 hrs. Most compounds significantly reduced the expression of all or most (4 to 5) genes assayed, except CUDC-907 (CUD) reduced 3 genes, ARV-825 (ARV) reduced *IFIT1* while increasing *ISG15*, prednisolone (Pred) increased *ISG15* and *MX1*, and dinaciclib (Dina) increased *HLA-C, ISG15*, and *MX1*. Echi – echinomycin, Givi – givinostat, Pano – panobinostat, Nemo – nemorubicin, Trip - triptonide. Bars represent mean ± standard deviation of duplicate PCR reactions from a single treated well. Statistical analysis by two-way ANOVA with Dunnett’s multiple comparisons to IFN-β treated sample. 4 – p<.05 for *HLA-C, IFIT1, ISG15*, and *MX1*. 5-p<.05 for all 5 genes assayed. Black or red asterisk – down- or up-regulation p<.05, respectively.

## Discussion

To expand the arsenal of therapeutics that might be effective for managing myositis, a high throughput screening campaign was performed to identify compounds that inhibit the type I IFN – MHC class I pathway in skeletal muscle. We report here the creation of a CRISPR/Cas9 genome-engineered myoblast in which a HiBit reporter was fused to endogenous HLA-B and was used to screen >4,500 compounds across titrations spanning several orders of magnitude. Active compounds were followed-up by HLA-ABC IF, RT-qPCR, and Western blotting to validate those that inhibit MHC class I transcription and filter out artifacts (**Supplementary Figure 6**). Through this series of assays, 57 compounds were active by HLA-B HiBit, HLA-ABC IF, and *HLA-B* RT-qPCR. Mechanism of action for 11 or 12 representative active compounds was investigated by Western blot and RT-qPCR, respectively. A summary of the numbers of compounds tested by each assay is given in **Table 1**. Summary statistics for efficacies, potencies, and cytotoxicity of 12 representative actives are shown in **Tables 2** and **3**, and molecular structures are shown in **Figure 6**. Summary statistics for all 62 compounds that were assayed by HLA-B HiBit, CTG, HLA-ABC IF, and HLA-B RT-qPCR are shown in **Supplementary Tables 5-9**. Classes of compounds that were active through all assays were inhibitors for: kinases, HDAC, DNA topoisomerase II, transcription factors, and glucocorticoids. Compounds that were active at decreasing MHC class I protein but far less effective on mRNA were inhibitors for: BRD4, proteasome, HSP90, Na+/K+ ATPase, and SERCA. Na+/K+ ATPase and SERCA are important for endoplasmic reticulum function (Patel, 2016; Primeau et al., 2018), through which nascent MHC class I protein is folded and processed, so inhibiting their function appears to decrease MHC class I protein production without affecting its transcription.

**Figure 6.**
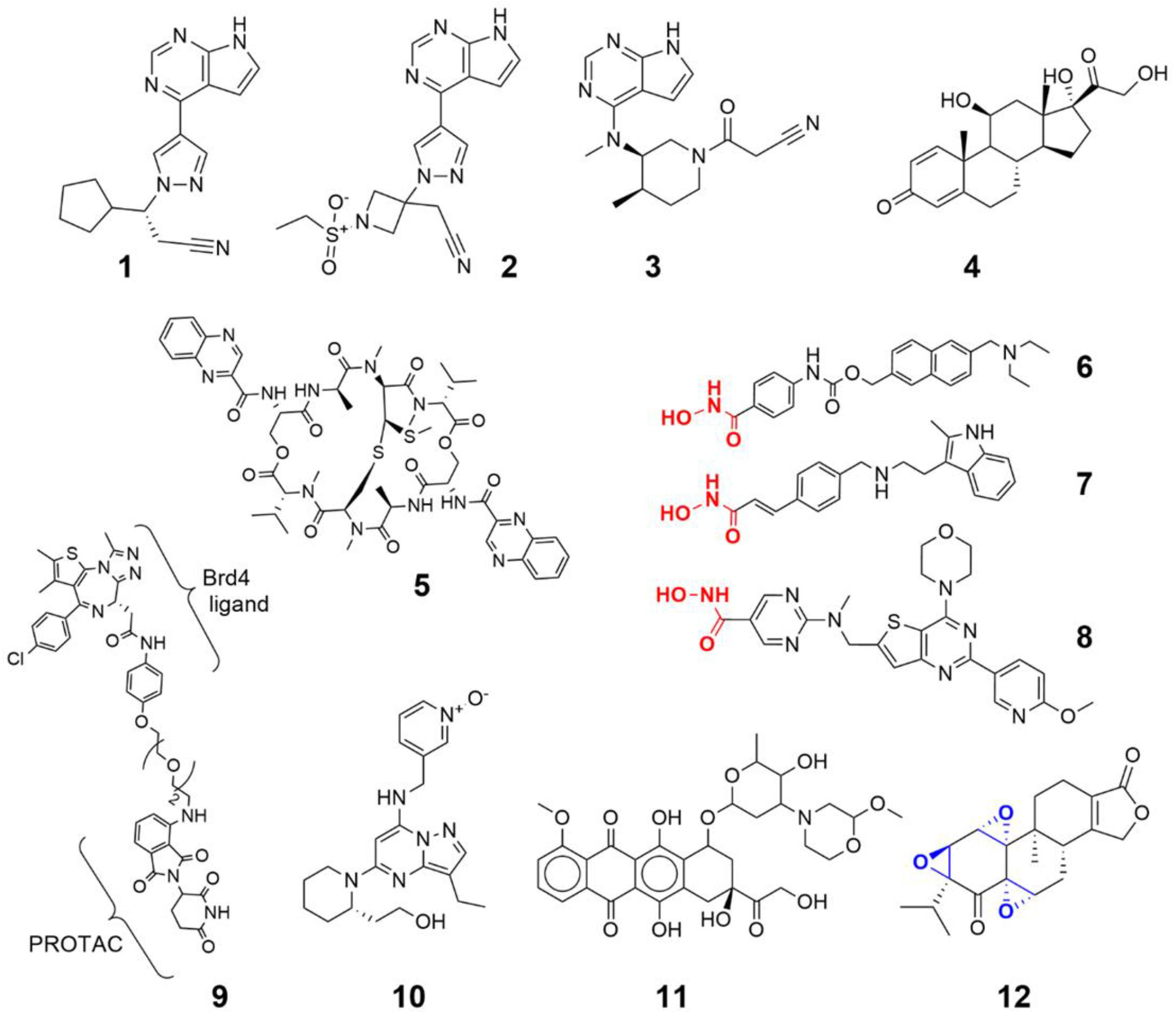
Chemical structures for representative active compounds. The JAK inhibitors ruxolitinib (**1**), baricitinib (**2**), and tofacitinib (**3**) have very similar core structures and similar activities for inhibiting IFN-β signaling. All the kinase inhibitors, including the PI3K/HDAC inhibitor CUDC-907 (**8**), and CDK1/2 inhibitor dinaciclib (**10)**, share a purine-like core, but differ in selectivity for the kinases they inhibit due to variant appendages. Givinostat (**6**) and panobinostat (**7**) are both HDAC inhibitors that have similar linear structures, containing a terminal hydroxamic acid (red), with panobinostat being 45x more potent than givinostat at inhibiting IFN-β-MHC class I pathway. The dual PI3K/HDAC inhibitor CUDC-907 also shares this hydroxamic acid moiety. Prednisolone (**4**) has a classic steroid structure and has high potency but very low efficacy in these assays. The HIF-1 inhibitor/DNA intercalator echinomycin (**5**) is a macrocyclic peptide ring. The BRD4 inhibitor ARV-825 (**9**) contains a proteolysis targeting chimera (PROTAC) linker to target BRD4 for ubiquitination and degradation. DNA topoisomerase II inhibitor nemorubicin (11) contains the anthracycline core. Although XPB transcription factor inhibitor triptonide (12) has a steroid structure, its activity is due to the reactive epoxides (blue) covalently binding to XPB.

**Table 1.** Summary of small molecule screening

| Screen | Assay | Wells per plate | Compounds tested | Titrations | % of total |
| --- | --- | --- | --- | --- | --- |
| Primary | HLA-B HiBit | 1536 | 6,576<br>(4,679 unique) | 7-11 | 100% |
| Counter | Retest HLA-B HiBit<br>& Cytotoxicity (CTG) | 1536 | 650 | 12 | 14% |
| Secondary | HLA-ABC<br>Immunofluorescence<br>& Cytotoxicity (Cell Count) | 1536 | 216 | 11 | 5% |
| Tertiary | HLA-B RT-qPCR | 384 | 62 | 6 | 1% |
| Mechanism | Multi Gene RT-qPCR | 384 | 12 | 1 | 0.3% |
| Mechanism | Western Blot | 6 | 11 | 1 | 0.2% |

CTG - CellTiter-Glo

**Table 2.**
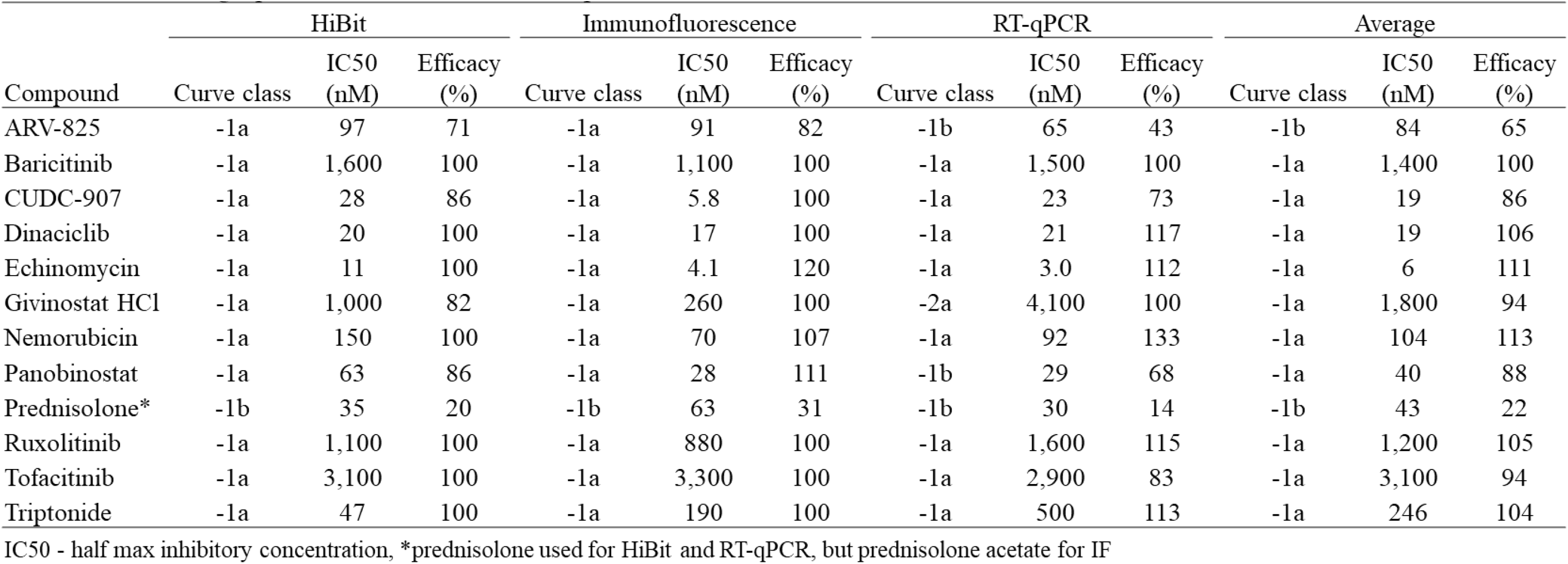
Pha1111acolo gic 2arameters for select active compounds

**Table 3.** Cytotoxicity parameters for select active compounds

| Compound | CTG |  |  | Cell count |  |  |
| --- | --- | --- | --- | --- | --- | --- |
|  | Curve class | IC50 (nM) | Efficacy (%) | Curve class | IC50 (nM) | Efficacy (%) |
| ARV-825 | -1b | 53 | 8 | 4 |  |  |
| Baricitinib | 4 |  |  | 4 |  |  |
| CUDC-907 | -1b | 3,500 | 40 | -3 |  | (10) |
| Dinaciclib | 4 |  |  | 4 |  |  |
| Echinomycin | 4 |  |  | 4 |  |  |
| Givinostat HCl | -3 |  | (18) | -3 |  | (25) |
| Nemorubicin | 4 |  |  | 4 |  |  |
| Panobinostat | -3 |  | (25) | 4 |  |  |
| Prednisolone* | 4 |  |  | 4 |  |  |
| Ruxolitinib | 4 |  |  | 4 |  |  |
| Tofacitinib | 4 |  |  | 4 |  |  |
| Triptonide | 4 |  |  | 4 |  |  |

CTG - CellTiter-Glo, IC50 - half max inhibitory concentration, \*prednisolone used for HiBit and RT-qPCR, but prednisolone acetate for IF, ( ) - efficacy at highest concentration

The quickest and most efficient approach to identifying therapies for a rare disease such as myositis is drug repurposing, which involves recognizing drugs already approved for other indications show efficacy in new indications (Pushpakom et al., 2018). The three libraries screened here were annotated, with available information on targets, mechanisms, pharmacokinetics, etc. Many of these compounds are approved by the US Food and Drug Administration (FDA), European Medicines Agency (EMA), or the Japanese Pharmaceuticals and Medical Devices Agency (PMDA) (Huang et al., 2011). With this annotated information, broad classes of compounds and targets can be identified. We can also use these libraries to investigate novel mechanisms of approved drugs. Looking through the primary screening data, none of the small molecule drugs currently used or tested in the clinic to treat myositis block the type I IFN – MHC class I pathway except JAK inhibitors and to a lesser extent glucocorticoids (**Supplementary Table 10**). Not only are these annotated libraries useful for efficient translation of therapies, but also for understanding basic biology using chemical probes to interrogate pathways. For example, our data showed HDAC inhibitors reduced IFN-β-stimulated MHC class I expression, indicating that HDACs play a role in this pathway not previously reported.

Successful high throughput screening impinges upon a rapid yet robust primary screen that can be miniaturized into 1536-well plates and retain biological relevance (Inglese et al., 2007a). For this purpose, we utilized the pro-luminescent reporter fragment HiBit, which is only 11 amino acids and encoded by 33 base-pairs, making it efficient for insertion into the genome and minimizes deleterious effects to the function and localization of its fusion protein (Schwinn et al., 2018). HiBit is a 1.3 kDa fragment of the luminescent NanoLuc, and when complexed with the larger 18 kDa portion of the enzyme (LgBit), they reconstitute a functional reporter to produce light in the presence of furimazine substrate. CRISPR/Cas9 is a sequence specific endonuclease by which pre-determined sequences of DNA can be cut with the use of synthetic gRNAs. Then, the endogenous DNA repair mechanism homology-directed repair inserts an exogenous DNA fragment flanked by homology arms (Jiang and Doudna, 2017; Ran et al., 2013). Here, HiBit was fused to the HLA-B*08:01 allele because it is the MHC class I allele within the ancestral 8.1 haplotype that is associated with myositis (Miller et al., 2015).

The largest class of active compounds through all assays were kinase inhibitors (26 kinase inhibitors / 57 actives). Half of them, 13, were JAK inhibitors, of which ruxolitinib, baricitinib, and tofacitinib are approved by the FDA and peficitinib is approved by the PMDA. The recently FDA approved upadacitinib was not in our libraries. Of the FDA approved JAK inhibitors in our libraries, all had about 100% efficacy, but ruxolitinib was the most potent, followed by baricitinib, and then tofacitinib (HiBit assay IC50 = 1100, 1600, 3100 nM, respectively). These results are in line with their relative potencies in published cell-free JAK1 assays where ruxolitinib is the most potent, followed by baricitinib, and then tofacitinib (IC50 = 3.3, 5.9, 112 nM respectively) (Changelian et al., 2003; Fridman et al., 2010; Quintás-Cardama et al., 2010). IFNAR transduces type I IFN signal via JAK1 and Tyk2. An important note is that tofacitinib has selectivity for JAK3 (IC50 = 1 nM), which is not involved in the IFN pathway, but is involved in transducing signals from cytokine receptors using the common γ chain, such as for IL-2, -4, -7, -9, -15, -21 (Changelian et al., 2003). So ruxolitinib or baricitinib have more selectivity and are more potent for IFN signaling, whereas tofacitinib produces more general immunosuppression by inhibiting a broader population of cytokines.

The other 13 active kinase inhibitors are investigational agents with various reported targets, including c-Jun N-terminal kinase (JNK), p21-activated kinase 4 (PAK4), PDZ-binding kinase, maternal embryonic leucine zipper kinase (MELK), TRAF2 and NCK-interacting protein kinase (TNIK), epidermal growth factor receptor (EGFR), inositol-requiring enzyme 1 (IRE1), protein kinase B (AKT), PI3K, and CDK. **Figure 3** showed CRC data for dinaciclib, a CDK1/2 inhibitor (Parry et al., 2010), and CUDC-907, a dual PI3K/HDAC inhibitor (Qian et al., 2012). Among the actives were two other compounds targeting CDK1/2, CGP-60474 and RGB-286147. Interestingly, the hill slope for the CDK1/2 inhibitors were very steep (>1), showing an ‘ultrasensitive’ response, which leads to a binary on-off type effect (Ferrell, 1996). This phenomenon can be the result of a variety of complex interactions within the cell and has been documented in CDK activation to regulate the cell cycle, as the entry and exit from cell division is an all-or-none response (Huang and Ferrell, 1996). A search of all screening data revealed that three FDA approved CDK4/6 inhibitors were either cytotoxic or inactive, indicating that perhaps CDK1/2, but not CDK4/6, are involved in this cell line or pathway, or the compounds have off-target effects, as kinase inhibitors can be promiscuous (Davis et al., 2011). The various identified kinase targets require further validation, which could be investigated by gene-targeted knock down with small interfering RNA (siRNA). As mentioned above, CUDC-907 is reported to inhibit both PI3K and HDAC. However, there was only one other active compound targeting PI3K (PIK-75), which also inhibits DNA-dependent protein kinase, while four FDA approved PI3K inhibitors had various effects ranging from cytotoxic, low potency, or low efficacy, providing little corroborating data to suggest PI3K inhibition affects this pathway. Alternatively, the effects of CUDC-907 might be due to its secondary HDAC inhibiting activity, as this class of compounds were active (discussed below) and share the aryl hydroxamic acid moiety (**Figure 6**).

Epigenetic and transcriptional modulators constitute another broad class of actives, including inhibitors of HDAC, DNA topoisomerase II, and transcription factors. Two FDA approved HDAC inhibitors were identified (panobinostat and vorinostat) as well as three investigational HDAC inhibitors (dacinostat, givinostat, and quisinostat). HDACs are not only involved in chromatin structure via removing acetyl groups from histones, but also known to regulate NF-κB and inflammation (Blanchard and Chipoy, 2005). Early trials in Duchenne muscular dystrophy patients and a mouse model showed improvement in muscle histology and reduced inflammation through the use of the HDAC inhibitor givinostat (Bettica et al., 2016; Consalvi et al., 2013). It would be valuable to extend these results to myositis in animal models or clinical trials. Three DNA topoisomerase II inhibitors were active: doxorubicin (FDA approved), nemorubicin (EMA approved), and aldoxorubicin (investigational). DNA topoisomerase II is important for unwinding DNA during both replication and transcription, so inhibiting this enzyme can reduce gene expression (Pommier et al., 2010). The hill slope for DNA topoisomerase II inhibitors were ultrasensitive. This may be explained here by the cooperative nature of transcriptional machinery binding to the DNA (Chu et al., 2009). The most potent and effective compound in all of the screening was echinomycin (HiBit assay efficacy = 100% and IC50 = 11 nM), which also exhibited an ultrasensitive response. Echinomycin is a peptide antibiotic that has been shown to intercalate DNA in a sequence-specific manner and inhibit hypoxia-inducible factor-1 (HIF-1) (Kong et al., 2005). There are reports that NF-κB can regulate HIF-1 (Van Uden et al., 2008), which may tie together the inflammatory and hypoxia pathways. This mechanism warrants further exploration, especially as hypoxia and HIF-1 are thought to play a role in myositis muscle pathology (Kinder et al., 2013). Triptonide and its analogue triptolide were actives that are reported to inhibit xeroderma pigmentosum type B (XPB) subunit of the general transcription factor TFIIH through covalent bonding between triptolide epoxides and a cysteine residue of XPB (He et al., 2015; Titov et al., 2011). Triptonide and triptolide are natural products isolated from the thunder god vine and used in ancient Chinese medicine for inflammatory diseases including arthritis (Qiu and Kao, 2003). These compounds displayed ultrasensitive responses in our assays, likely due to their inhibition of transcription machinery as described above. Inhibition of the type I IFN – MHC class I pathway may be one mechanism by which triptonide is anti-inflammatory. Finally, four investigational NF-κB inhibitors were active: withaferin A, parthenolide, CBL-0137, and TPCA-1, which was expected as NF-κB is a known mediator of the IFN pathway (Pfeffer, 2011).

The current first-line therapy for dermatomyositis, polymyositis, and necrotizing autoimmune myopathy is prednisone (Dalakas, 2015). However, it has serious side effects including metabolic and endocrine disturbances, osteopenia and bone breaks, psychiatric changes, skeletal muscle atrophy and weakness, hypertension, and renal disturbances (Hoffman et al., 2018). These are especially concerning in children, who can exhibit stunting of growth, delay in puberty, weight gain, mood disturbances, and fragile bones. From our data, glucocorticoids had very low efficacy at inhibiting type I IFN – MHC class I pathway (about 20%), and increased expression of several ISGs in the presence of IFN-β. Lundberg et al. have observed residual MHC class I expression in muscle and refractory weakness in some patients treated with prednisolone, despite elimination of infiltrating inflammatory cells (Lundberg et al., 2000). This indicates that eliminating infiltrating inflammatory cells through glucocorticoid treatment often doesn’t cure muscle weakness or stop inflammatory signals leading to MHC class I expression. Therefore, there is a need for more targeted therapies to eliminate the refractory muscle weakness and inflammation.

This study is the first to pursue an HTS approach to repurpose therapies for myositis. Our use of CRISPR/Cas9 genome-engineering to insert a small reporter into the endogenous locus of *HLA-B* extents our work on using genome-editing to develop HTS reporter cell lines by encompassing post-translational responses to compound treatment (Hasson et al., 2015; Inglese et al., 2014). The assays reported here also advance the field of HTS with our utilization of cells from the target tissue type (myoblasts from skeletal muscle), rather than the often used HEK293 or HeLa cells. Reporter integration at the physiological cell type genomic locus is particularly important for the mechanism of action interpretation of compounds exerting chromatin modifying effects, where genome context is lost in randomly integrated reporter systems. Using myoblasts, we may identify mechanisms that are unique to this cell type. These assays could be further utilized to screen diverse/novel chemical matter for new chemical probes, investigate the regulation of MHC class I expression in muscle, and test for inflammatory triggers that may precipitate myositis. Compared to traditional single-point HTS, our titration-based qHTS approach has the advantage of profiling the full pharmacologic response to compounds by showing the efficacy, potency, and hill slope (Inglese et al., 2006). With this information we can compare which drugs are most potent in a class, like ruxolitinib as the most potent JAK inhibitor, and glean some mechanistic information from hill slopes, like ultrasensitiviy of inhibitors for DNA topoisomerase II, transcription factors, and CDKs.

The active agents identified here might have broader applications to treat other autoimmune conditions that also have a type I IFN response, such as systemic lupus erythematosus, Sjögrens syndrome, systemic sclerosis, and rheumatoid arthritis (Rönnblom, 2016). Compounds that inhibit MHC class I expression could have therapeutic value in autoimmunity associated with MHC class I misfolding, as is seen in HLA-B27-associated ankylosing spondylitis (Colbert et al., 2014).

Limitations of this work include the presumption that the type I IFN – MHC class I pathway is pathogenic in myositis. It may be that this pathway, or components of it, have differing roles in different subtypes of myositis. For example, the type I IFN signature is most prevalent in dermatomyositis (Gallay et al., 2019). It may also be that type I IFN or MHC class I expression are consequences rather than causes of pathology in humans. The exact targets and mechanisms of some of the compounds reported here are not completely understood and should be investigated further. Finally, these results reflect a simplified *in vitro* system, where safety and efficacies need to be confirmed in animal models and clinical trials before translation to patients.

## Acknowledgments

We would like to thank NCATS staff with compound management, automation, and informatics: Z. Itkin, J. Travers, S. Frebert, C. Klumpp-Thomas, Y. Wang, J. Braisted, and B. Queme. K. Cheng, PhD (NCATS) was helpful in planning the CRISPR/Cas9 genome-engineering. We thank A. Dutra, Ph.D. of NHGRI for karyotyping, and R. Pellegrino, PhD with L. Tian at CHOP for WGS. We are also grateful for consultation with members of the Cure JM medical advisory board K. Nagaraju, PhD, DVM (Binghamton University) and L. Rider, MD (National Institute of Environmental Health Sciences). We are appreciative for research materials including C24cl48 myoblasts from V. Mouly, PhD (Institut de Myologie) and HC-10 antibody from R.A. Colbert, PhD (National Institute of Arthritis and Musculoskeletal and Skin Diseases). We thank D. Leja (NHGRI) for Figure 1 illustrations.

---

**Supplementary Figure 1.**
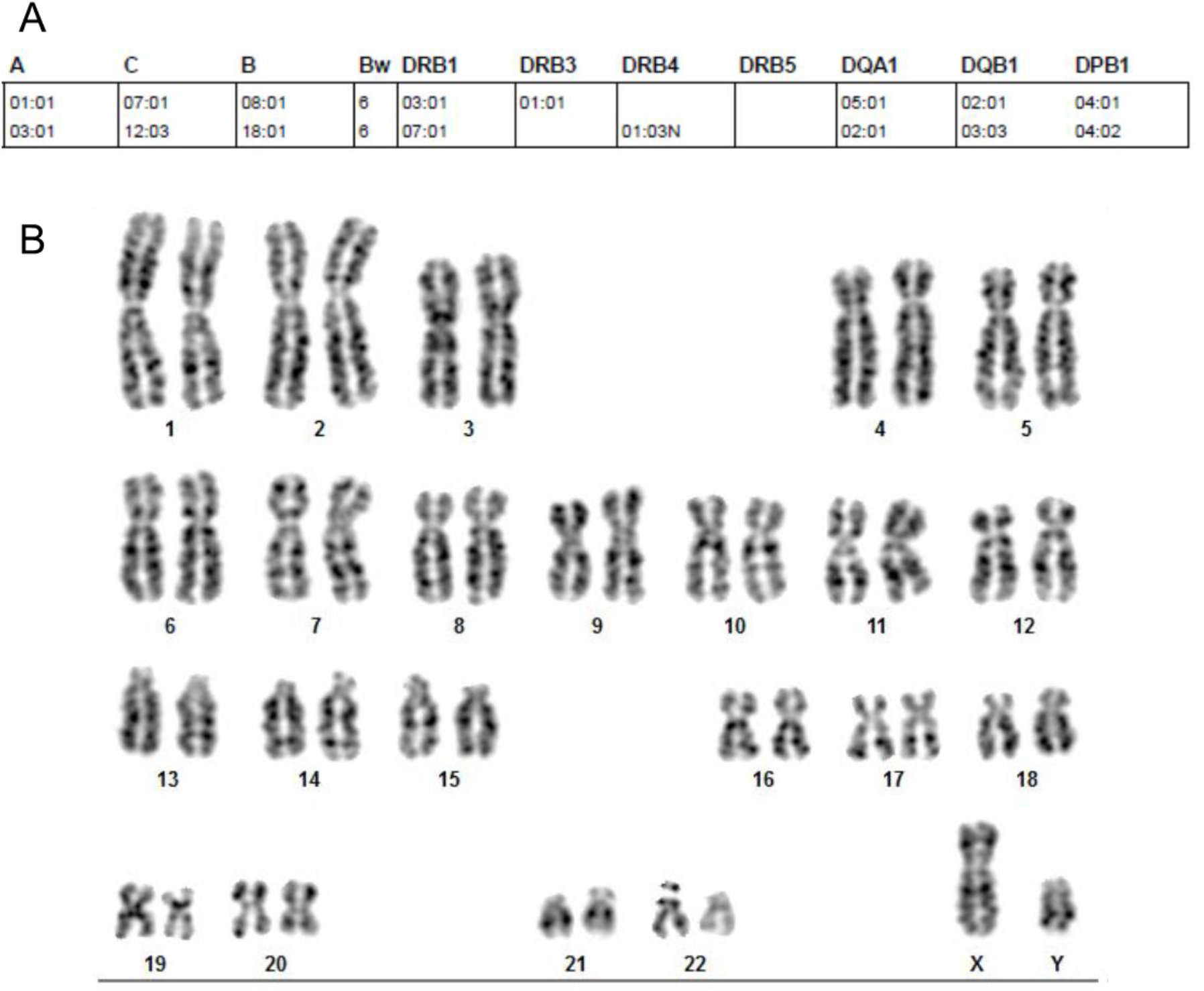
HLA genotyping and karyotyping for immortalized human myoblasts C25cl48. **A,** DNA was extracted from C25cl48 cells and HLA genotypes determined by next generation sequencing, high resolution HLA typing. C25cl48 cells were determined to be heterozygous at the *HLA-B* locus, and we targeted HLA-B*08:01 for genome-engineering. **B,** Karyotyping showed C25cl48 cells to be normal 46,XY.

**Supplementary Figure 2.**
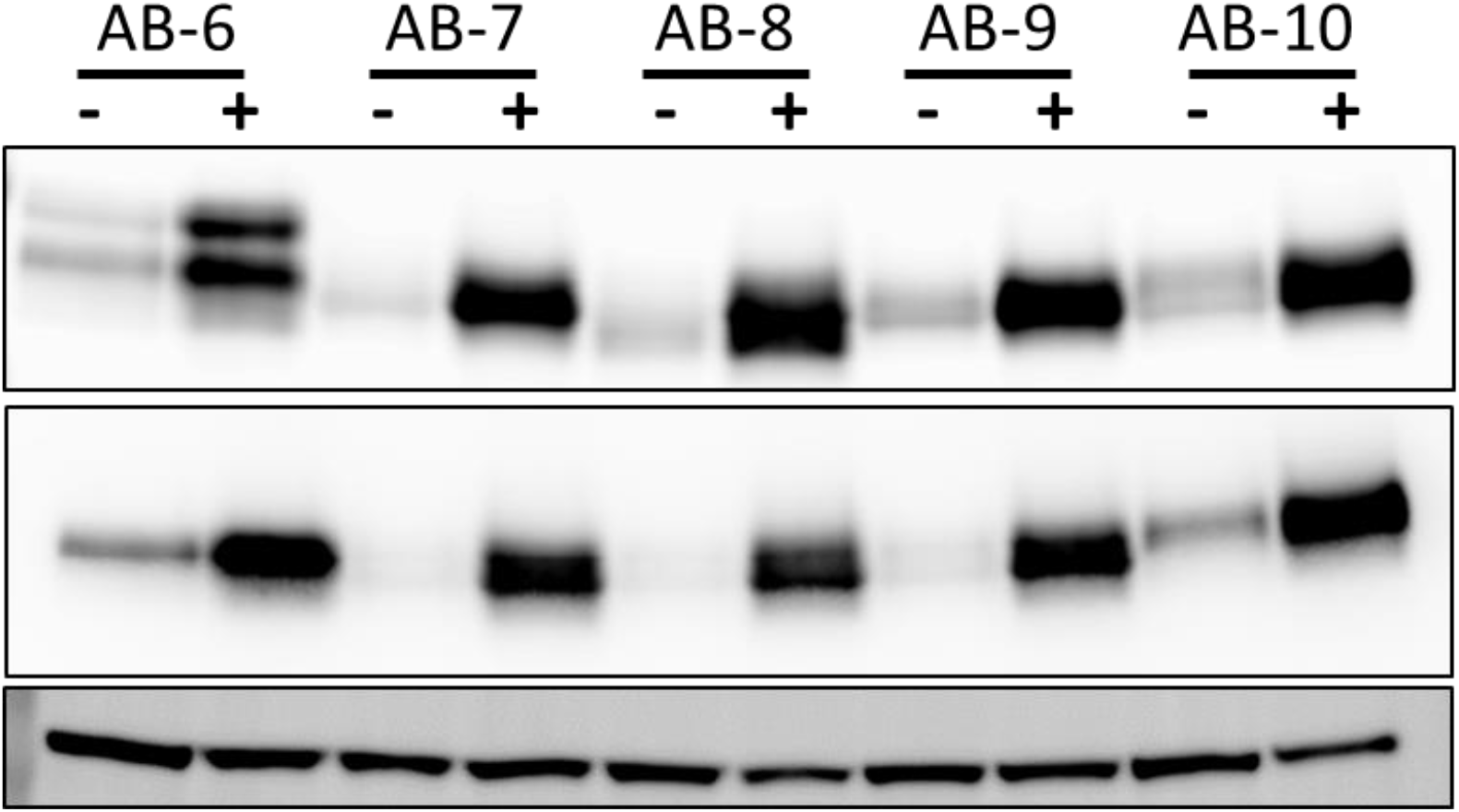
Western blot analysis of HLA-ABC, HiBit, and β-tubulin loading control in several additional genome-engineered clones treated + 8.0 ng/ml IFN-β for 24 hrs showed up-regulation of endogenous HLA-ABC and HiBit with IFN-β treatment.

**Supplementary Figure 3.**
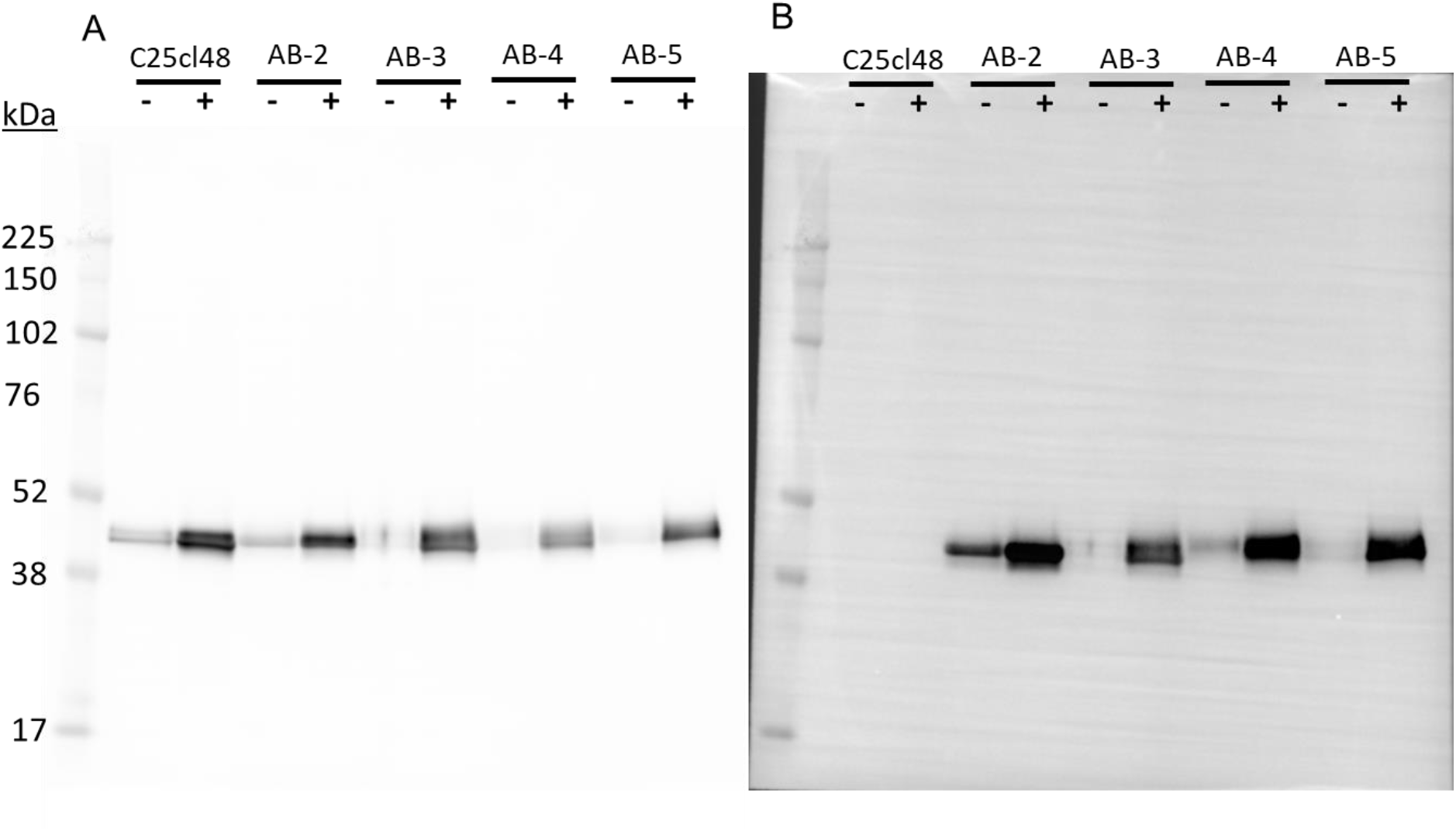
Full membrane images of Western blot analysis for HLA-ABC and HiBit in parental C25cl48 cells and several genome-engineered clones treated + 8.0 ng/ml IFN-β for 24 hrs showed the HiBit band migrated at the same molecular weight (∼45 kDa) and was up-regulated in the same manner as endogenous HLA-ABC, which provides evidence that the insertion was on-target. Also, HiBit signal was only detected at this one location, indicating there was no off-target tagging of other proteins. A, Membrane probed with HLA-ABC antibody. B, Membrane probed with LgBit and furimazine substrate.

**Supplementary Table 1.** Summary statistics for HLA-B HiBit primary screen, LOPAC1280 library

| Parameters | Assay Statistics |  |
| --- | --- | --- |
| 1536-well plates | 7 (+2 DMSO) |  |
| Compounds tested | 1,280 |  |
| Points per titration | 7 |  |
| Data points | 8,960 |  |
| Control condition | IFN-β (0.18 nM = 4.0 ng/ml) | |
| | DMSO | IFN-β |
| HiBit Luminescence | $537.7 \pm 86.7$ | $16,482.5 \pm 684.3$ |
| CV | $15.7 \pm 5.1$ | $4.1 \pm 0.7$ |
| S/B | $32.5 \pm 9.3$ | |
| Z' factor | $0.89 \pm 0.03$ | |

S/B - Signal/Background, CV - Coefficient of variation

**Supplemental Table 2.** HLA-B HiBit clones whole genome sequencing coverage

| Sample<br>ID | Average depth of<br>sequencing coverage |
| --- | --- |
| AB-3 | 38 |
| AB-5 | 38 |
| AB-7 | 31 |
| AB-11 | 42 |
| WT | 48 |

**Supplemental Table 3.**
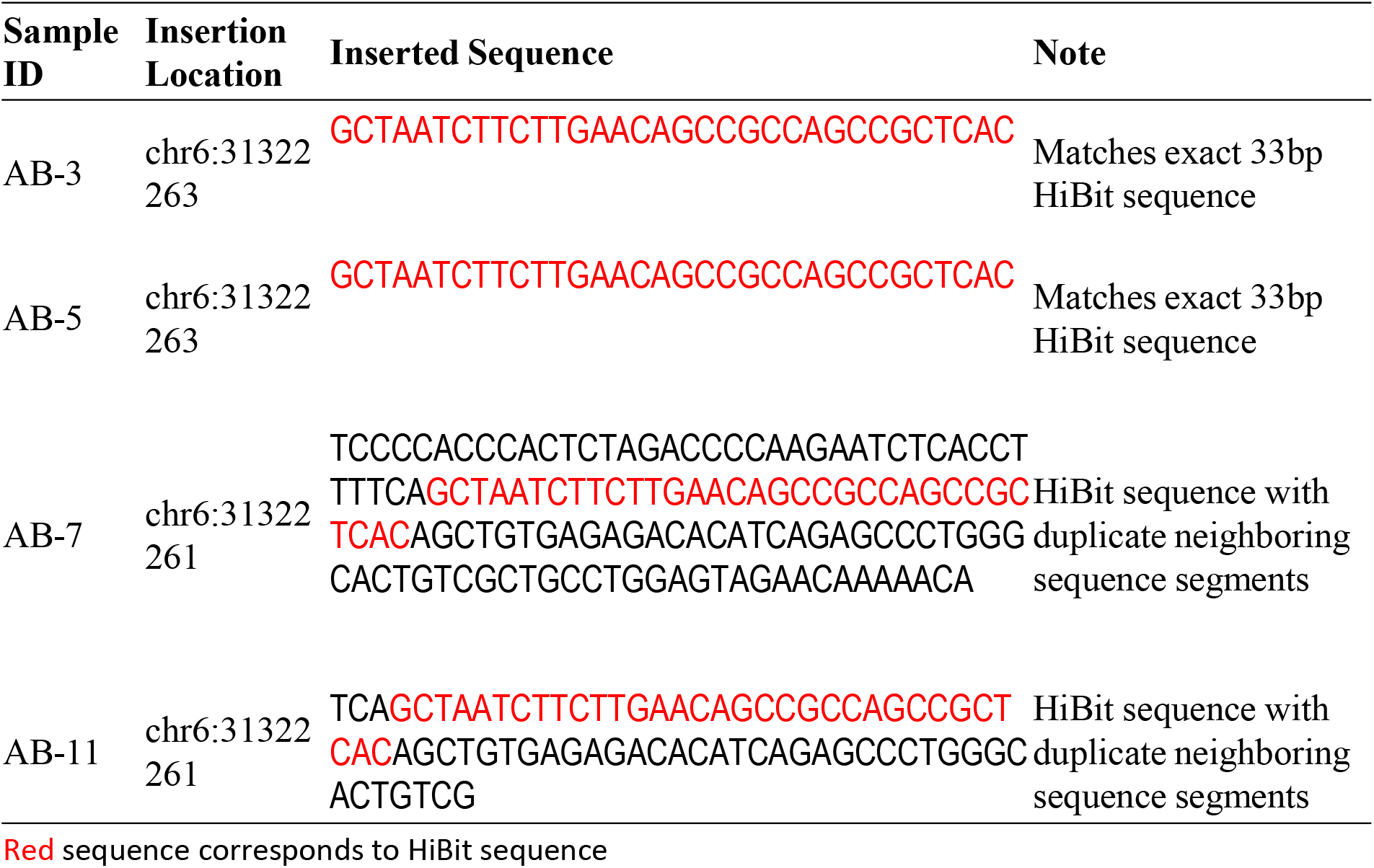
HLA-B HiBit clones whole genome sequencing insert analysis

**Supplementary Figure 4.**
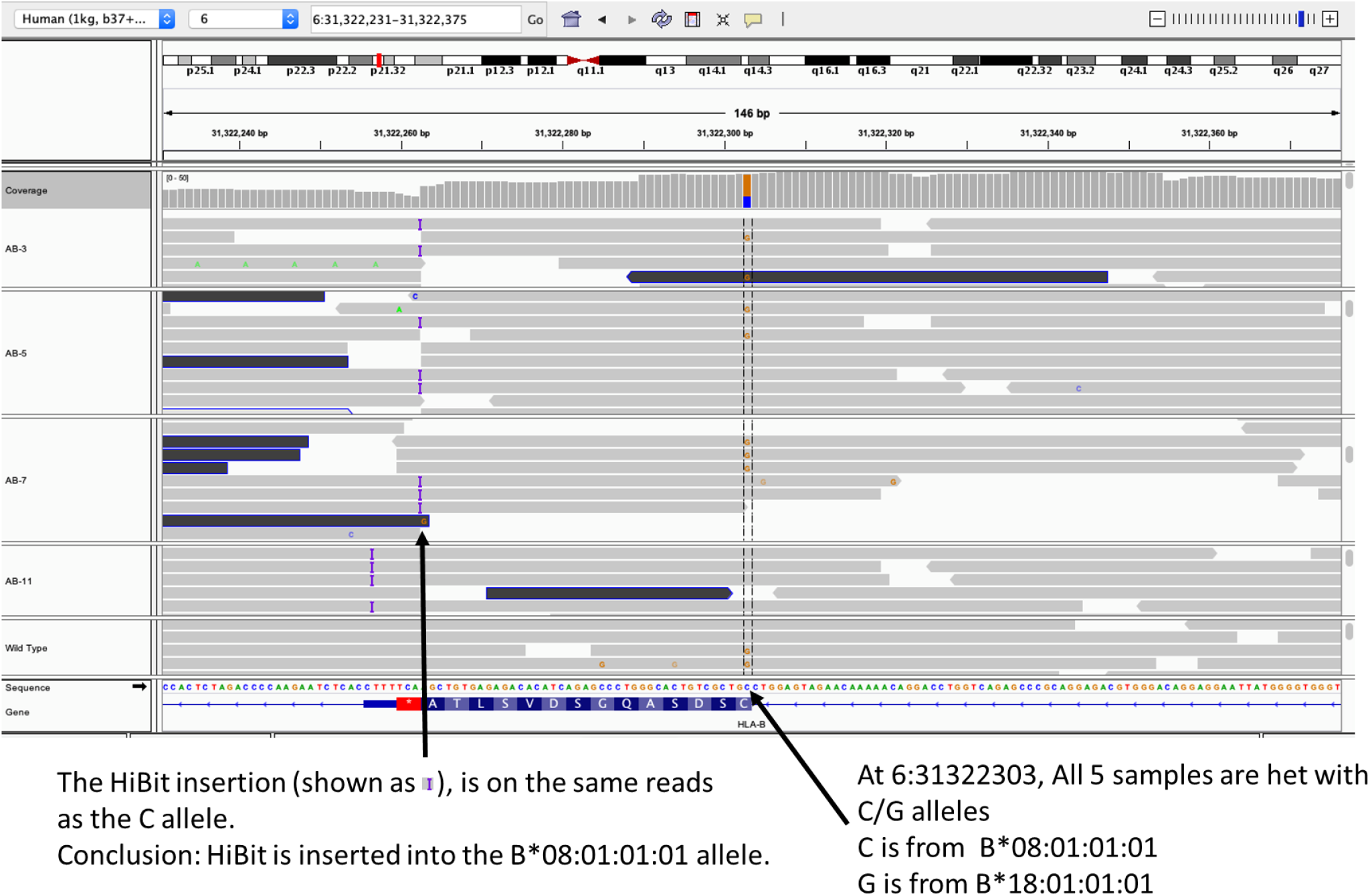
Phasing analysis of whole genome sequencing for HLA-B HiBit clones showed insertion at the desired HLA-B*08:01:01:01 but not at the HLA-B*18:01:01:01 allele for all four clones AB-3, -5, -7, and -11.

**Supplementary Table 4.** Primary libraries screened

| Library | Compounds | Titirations | Dilution Factor | Z'-factor |
| --- | --- | --- | --- | --- |
| LOPAC | 1,280 | 7 | 1:4.3 | 0.89 |
| NPC | 2,816 | 8 | 1:5 | 0.82 |
| MIPE5.0 | 2,480 | 11 | 1:3 | 0.80 |
| Sum | 6,576 |  |  |  |
| Unique | 4,679 |  |  |  |

LOPAC - Library of pharmacologically active compounds, NPC - NCATS pharmaceutical collection, MIPE5.0 - Mechanism interrogation plate version 5.0

**Supplementary Figure 5.**
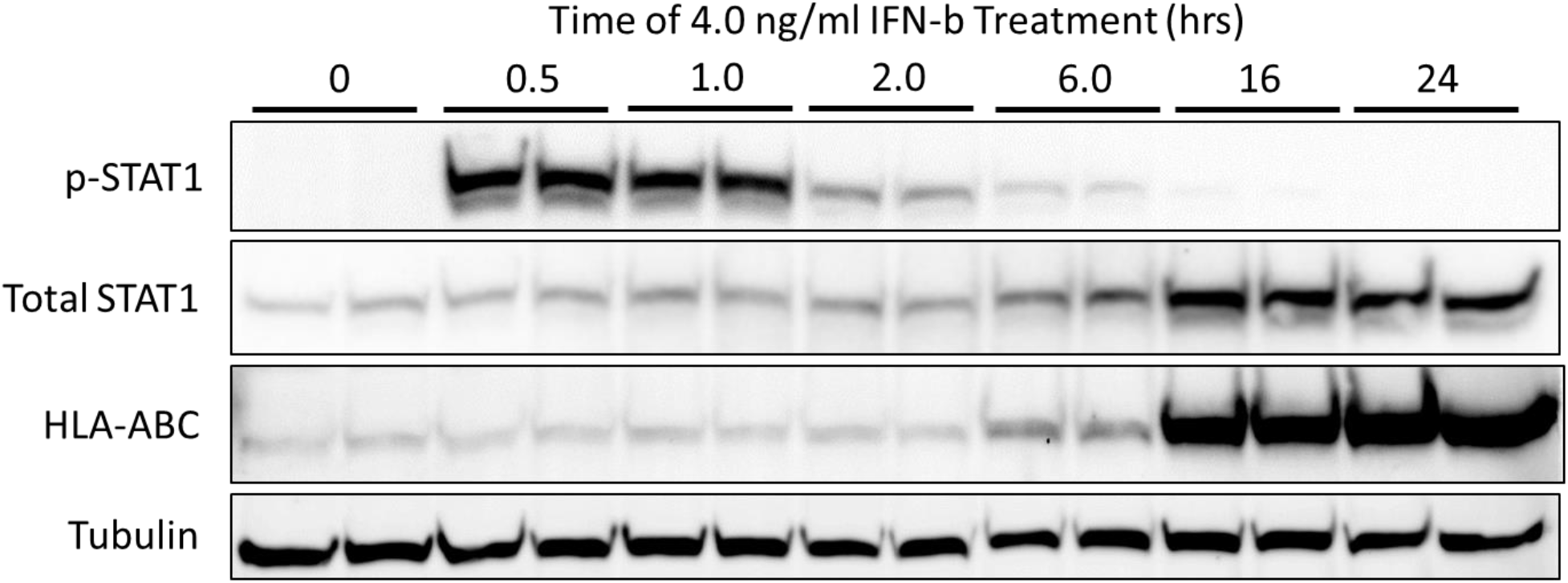
Western blot analysis of phospho-STAT1 (p-STAT1), total STAT1, HLA-ABC, and tubulin loading control for C25cl48 cells treated with 4.0 ng/ml IFN-β for 0.5, 1.0, 2.0, 6.0, 16 and 24 hrs. Stimulation with IFN-β caused phosphorylation of STAT1 by 0.5 hrs of treatment, began to reduce at 2.0 hrs, and was back to baseline by 24 hrs. Whereas stimulation with IFN-β caused total STAT1 and HLA-ABC to be up-regulated starting at 6.0 hrs and greatly increasing by 16 hrs. Duplicate adjacent lanes represent duplicate treated wells from 6 well plates.

**Supplementary Figure 6.**
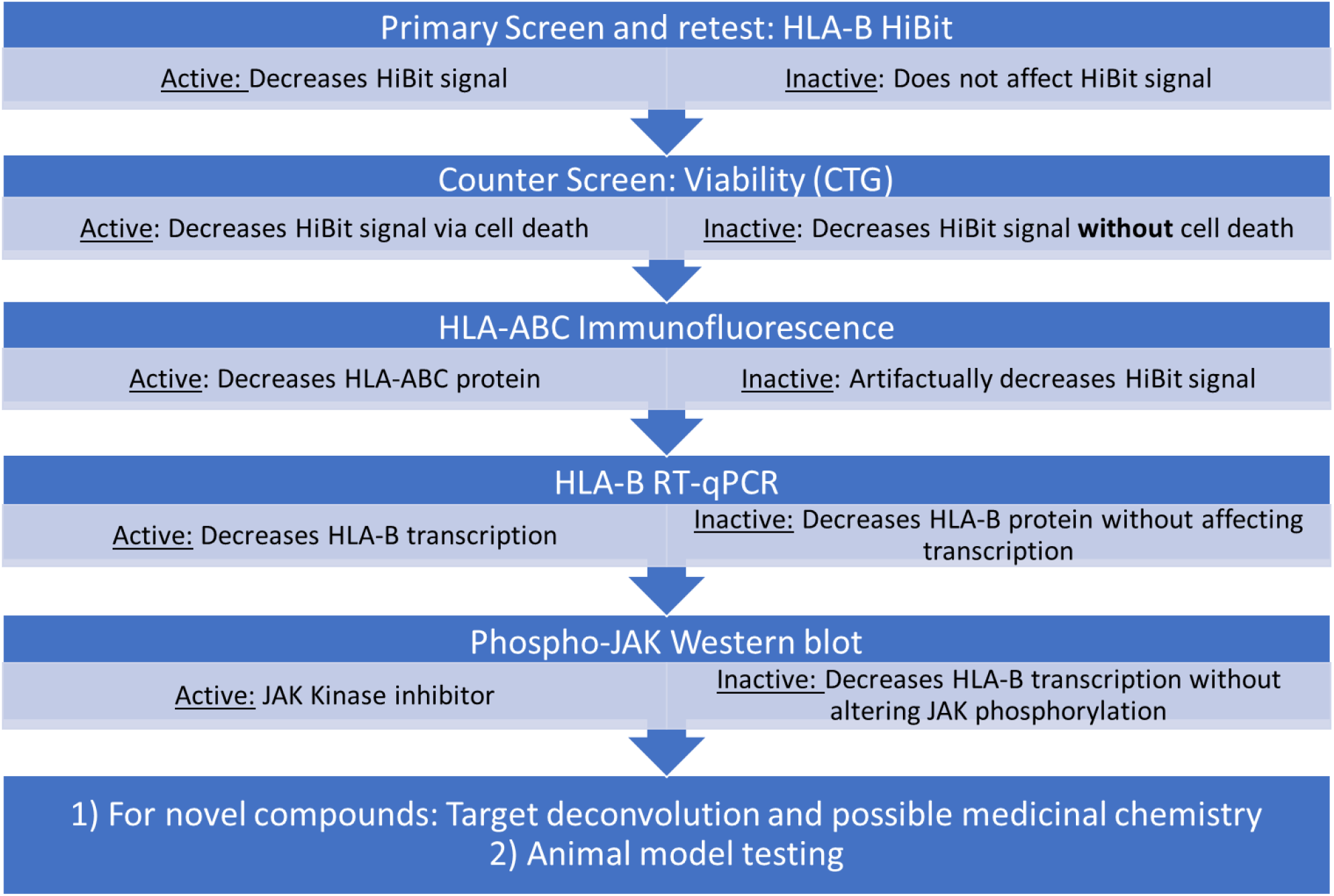
Flow chart showing the series of follow-up assays for screening compounds and inferences drawn from results at each stage. This strategy will reduce assay artifacts, ensure biological pharmacology of active compounds, and help elucidate mechanisms of action. Compounds identified in primary screening are retested by HLA-B HiBit assay as well as the cytotoxicity CTG assay to eliminate compounds that are cytotoxic. Minimally toxic actives are then assayed by IF of endogenous HLA-ABC protein in unedited C25cl48 cells to eliminate compounds that artifactually decrease HiBit signal. Next, actives are assayed by RT-qPCR, where inactivity at this stage represents compounds that decrease HLA-B protein without affecting transcription, and activity at this stage indicate inhibitors of HLA-B transcription. Testing for JAK phosphorylation shows compounds that inhibit transcription via inhibition of JAK activation or some other mechanism such as epigenetic regulation. Compounds with unknown targets or mechanisms need to be deconvoluted and medicinal chemistry can optimize leads. Finally, leads need to be tested in animal models to show safety and efficacy before clinical trials.

**Supplementary Table 5.**
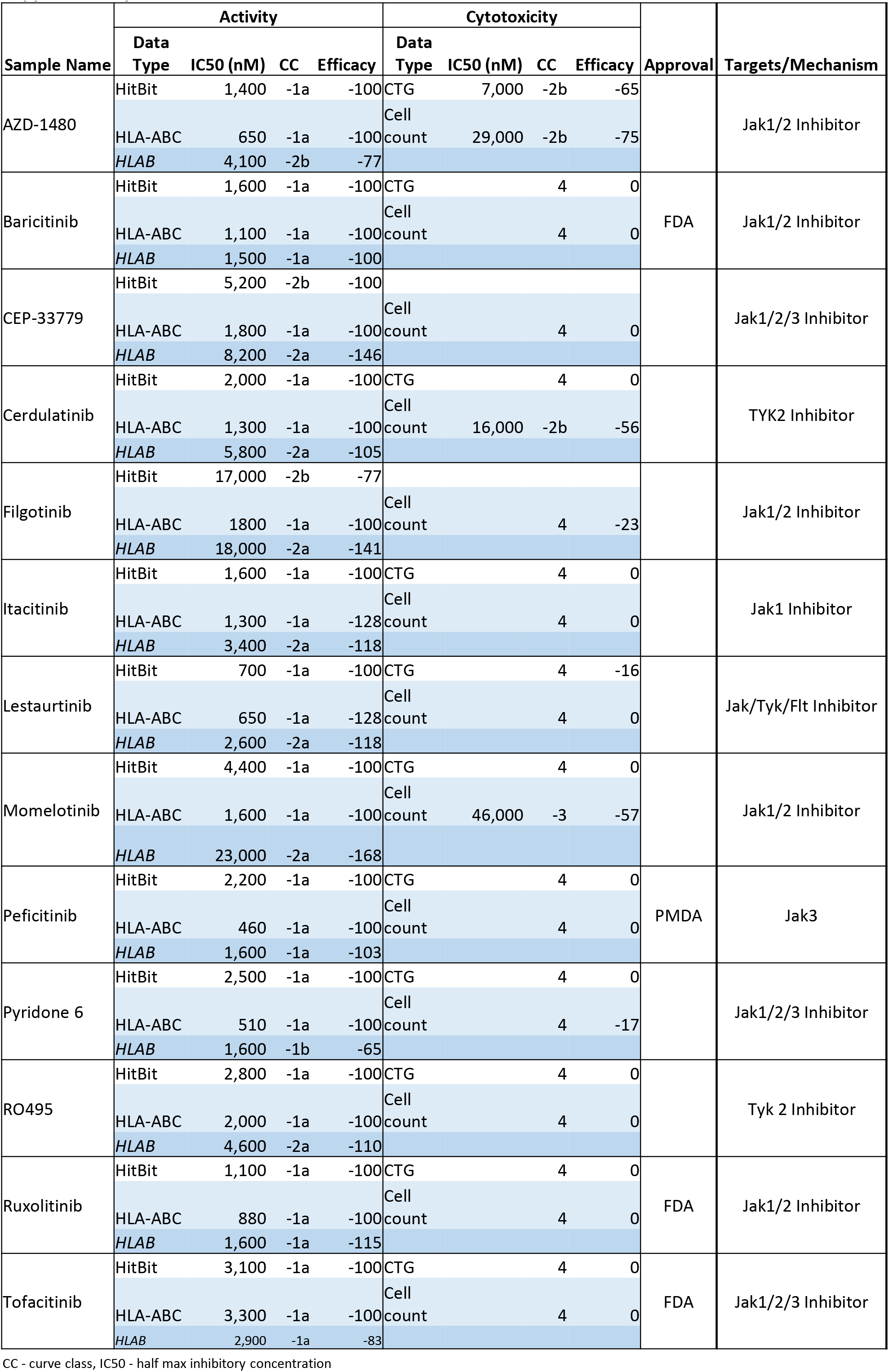
JAK inhibitors

**Supplementary Table 6.** Other kinase inhibitors

| Sample Name | Activity |  |  |  | Cytotoxicity |  |  |  | Approval | Targets/Mechanism |
| --- | --- | --- | --- | --- | --- | --- | --- | --- | --- | --- |
|  | Data Type | IC50 (nM) | CC | Efficacy | Data Type | IC50 (nM) | CC | Efficacy |  |  |
| A-443654 | HitBit | 1,100 | -1a | -100 | CTG | 4,400 | -2b | -37 |  | AKT Inhibitor |
|  | HLA-ABC | 230 | -1a | -83 | Cell count | 26,000 | -2a | -100 |  |  |
|  | HLAB | 210 | -1a | -109 |  |  |  |  |  |  |
| APY29 | HitBit | 1,400 | -1a | -100 | CTG | 2,000 | 1b | 25 |  | IRE1 Inhibitor |
|  | HLA-ABC | 650 | -1a | -100 | Cell count | 23,000 | -2b | -49 |  |  |
|  | HLAB | 650 | -1a | -121 |  |  |  |  |  |  |
| CGP-60474 | HitBit | 130 | -1a | -100 | CTG |  | 4 | 0 |  | CDK1/2, SMG1, mTOR Inhibitor |
|  | HLA-ABC | 81 | -1a | -100 | Cell count |  | 4 | 0 |  |  |
|  | HLAB | 120 | -1a | -121 |  |  |  |  |  |  |
| CUDC-101 | HitBit | 2,500 | -1a | -100 | CTG |  | 4 | 0 |  | EGFR, HDAC Inhibitor |
|  | HLA-ABC | 290 | -1a | -100 | Cell count |  | 4 | 0 |  |  |
|  | HLAB | 410 | -1b | -68 |  |  |  |  |  |  |
| Dinaciclib (SCH727965) | HitBit | 20 | -1a | -100 | CTG |  | 4 | 0 |  | CDK1/2/5/9 Inhibitor |
|  | HLA-ABC | 17 | -1a | -100 | Cell count |  | 4 | 0 |  |  |
|  | HLAB | 21 | -1a | -117 |  |  |  |  |  |  |
| NCB-1026 | HitBit | 560 | -1a | -100 | CTG |  | 4 | 0 |  | TNIK Inhibitor, Wnt Signaling Inhibitor |
|  | HLA-ABC | 1,000 | -1a | -128 | Cell count |  | 4 | 0 |  |  |
|  | HLAB | 650 | -1a | -136 |  |  |  |  |  |  |
| OTS-167 | HitBit | 99 | -1a | -100 | CTG |  | 4 | -15 |  | MELK Inhibitor |
|  | HLA-ABC | 46 | -1a | -118 | Cell count | 15,000 | -2a | -100 |  |  |
|  | HLAB | 120 | -1a | -130 |  |  |  |  |  |  |
| OTS964 | HitBit | 79 | -1a | -100 | CTG |  | 4 | 0 |  | PDZ-binding kinase Inhibitor |
|  | HLA-ABC | 81 | -1a | -100 | Cell count |  | 4 | 0 |  |  |
|  | HLAB | 52 | -1a | -136 |  |  |  |  |  |  |
| PF-3758309 | HitBit | 63 | -1a | -100 | CTG |  | 4 | 0 |  | PAK1/4 Inhibitor |
|  | HLA-ABC | 58 | -1a | -100 | Cell count |  | 4 | 0 |  |  |
|  | HLAB | 1,800 | -1a | -135 |  |  |  |  |  |  |
| PIK-75 | HitBit | 50 | -1a | -100 | CTG |  | 4 | 0 |  | CDK9, PI3K Inhibitor |
|  | HLA-ABC | 160 | -1a | -128 | Cell count |  | 4 | 0 |  |  |
|  | HLAB | 230 | -1a | -136 |  |  |  |  |  |  |
| RGB-286147 | HitBit | 35 | -1a | -100 | CTG |  | 4 | 26 |  | CDK1/2/3/7/9 Inhibitor |
|  | HLA-ABC | 23 | -1a | -100 | Cell count |  | 4 | 0 |  |  |
|  | HLAB | 37 | -1a | -90 |  |  |  |  |  |  |
| SR-3306 | HitBit | 2,500 | -1a | -100 | CTG |  | 4 | 0 |  | JNK 1/2/3 Inhibitor |
|  | HLA-ABC | 1,800 | -1a | -100 | Cell count |  | 4 | 0 |  |  |
|  | HLAB | 1,500 | -1b | -64 |  |  |  |  |  |  |

CC - curve class, IC50 - half max inhibitory concentration

**Supplementary Table 7.** Transcriptional and epigenetic modulators

| Sample Name | Activity |  |  |  | Cytotoxicity |  |  |  | Approval | Targets/Mechanism |
| --- | --- | --- | --- | --- | --- | --- | --- | --- | --- | --- |
|  | Data Type | IC50 (nM) | CC | Efficacy | Data Type | IC50 (nM) | CC | Efficacy |  |  |
| Aldoxorubicin hydrochlorid | HitBit | 2,800 | -1a | -100 | CTG |  | 4 | 0 |  | DNA topoisomerase II inhibitor (pro-Doxorubicin) |
|  | HLA-ABC | 1,000 | -1a | -100 | Cell count |  | 4 | 0 |  |  |
|  | HLAB | 4,600 | -1a | -143 |  |  |  |  |  |  |
| CBL-0137 | HitBit | 1,100 | -1a | -100 | CTG |  | 4 | 0 |  | NF-kappaB Activation Inhibitor |
|  | HLA-ABC | 650 | -1a | -100 | Cell count |  | 4 | 0 |  |  |
|  | HLAB | 1,800 | -1a | -136 |  |  |  |  |  |  |
| CUDC-907 | HitBit | 28 | -1a | -86 | CTG | 3,500 | -1b | -40 |  | PI3K/HDAC Inhibitor |
|  | HLA-ABC | 6 | -1a | -100 | Cell count |  | 4 | 0 |  |  |
|  | HLAB | 23 | -1b | -73 |  |  |  |  |  |  |
| Dacinostat | HitBit | 200 | -1a | -86 | CTG |  | 4 | 0 |  | HDAC Inhibitor |
|  | HLA-ABC | 36 | -1a | -100 | Cell count |  | 4 | 0 |  |  |
|  | HLAB | 120 | -1b | -55 |  |  |  |  |  |  |
| Doxorubicin | HitBit | 500 | -1a | -100 | CTG |  | 4 | -17 | FDA | DNA Topoisomerase II Inhibitor |
|  | HLA-ABC | 230 | -1a | -100 | Cell count |  | 4 | 0 |  |  |
|  | HLAB | 1,000 | -1a | -156 |  |  |  |  |  |  |
| Echinomycin | HitBit | 11 | -1a | -100 | CTG |  | 4 | 0 |  | HIF-1 seq specific DNA intercalater |
|  | HLA-ABC | 4 | -1a | -120 | Cell count |  | 4 | 0 |  |  |
|  | HLAB | 3 | -1a | -112 |  |  |  |  |  |  |
| Givinostat hydrochloride | HitBit | 1,000 | -1a | -82 | CTG |  | 4 | -30 |  | HDAC Inhibitor |
|  | HLA-ABC | 260 | -1a | -100 | Cell count | 51,000 | -3 | -52 |  |  |
|  | HLAB | 4,100 | -2a | -100 |  |  |  |  |  |  |
| Nemorubicin | HitBit | 150 | -1a | -100 | CTG |  | 4 | 0 | EMA | DNA Topoisomerase II Inhibitor |
|  | HLA-ABC | 70 | -1a | -107 | Cell count |  | 4 | 0 |  |  |
|  | HLAB | 92 | -1a | -133 |  |  |  |  |  |  |
| Panobinostat | HitBit | 63 | -1a | -86 | CTG |  | 4 | -100 | FDA | HDAC Inhibitor |
|  | HLA-ABC | 28 | -1a | -111 | Cell count |  | 4 | 0 |  |  |
|  | HLAB | 29 | -1b | -68 |  |  |  |  |  |  |
| Parthenolide | HitBit | 7,000 | -1a | -100 | CTG |  | 4 | 0 |  | Adiponectin receptor protein 2, HDAC, Microphthalmia-associated transcription factor inhibitor, NF-kappaB Activation Modulator |
|  | HLA-ABC | 7,300 | -1a | -100 | Cell count | 12,000 | -1a | -100 |  |  |
|  | HLAB | 16,000 | -2a | -126 |  |  |  |  |  |  |
| Quisinostat | HitBit | 160 | -1a | -100 | CTG | 1,400 | -1b | -30 |  | HDAC Inhibitor |
|  | HLA-ABC | 9 | -1a | -100 | Cell count |  | 4 | 0 |  |  |
|  | HLAB | 23 | -1b | -56 |  |  |  |  |  |  |
| TPCA-1 | HitBit | 1,800 | -1a | -100 | CTG |  | 4 | -38 |  | IKK beta Inhibitor |
|  | HLA-ABC | 1,400 | -1a | -121 | Cell count |  | 4 | 0 |  |  |
|  | HLAB | 10,000 | -2a | -117 |  |  |  |  |  |  |
| Triptonide | HitBit | 47 | -1a | -100 | CTG |  | 4 | 0 |  | XPB transcription factor inhibitor, Immunosuppressant |
|  | HLA-ABC | 190 | -1a | -100 | Cell count |  | 4 | 0 |  |  |
|  | HLAB | 500 | -1a | -113 |  |  |  |  |  |  |
| Vorinostat (SAHA) | HitBit | 3,500 | -1a | -100 | CTG |  | 4 | 0 | FDA | HDAC Inhibitor |
|  | HLA-ABC | 910 | -1a | -100 | Cell count |  | 4 | 0 |  |  |
|  | HLAB | 1,800 | -1b | -69 |  |  |  |  |  |  |
| Withaferin A | HitBit | 880 | -1a | -100 | CTG | 12,000 | -2b | -100 |  | Glu [NMDA] receptor, HSP90, PANC-1, NF-kappaB Activation Inhibitor |
|  | HLA-ABC | 730 | -1a | -100 | Cell count | 2,000 | -1a | -100 |  |  |
|  | HLAB | 1,300 | -1a | -85 |  |  |  |  |  |  |

CC - curve class, IC50 - half max inhibitory concentration

**Supplementary Table 8.** Other actives

| Sample Name | Activity |  |  |  | Cytotoxicity |  |  |  | Approval | Targets/Mechanism |
| --- | --- | --- | --- | --- | --- | --- | --- | --- | --- | --- |
|  | Data Type | IC50 (nM) | CC | Efficacy | Data Type | IC50 (nM) | CC | Efficacy |  |  |
| BMS-214662 | HitBit | 1,300 | -1a | -100 | CTG |  | 4 | -24 |  | Farnesyltransferase (geranylgeranyl pyrophosphate synthase) Inhibitor |
|  | HLA-ABC | 580 | -1a | -100 | Cell count | 20,000 | 2b | 45 |  |  |
|  | HLAB | 650 | -1b | -74 |  |  |  |  |  |  |
| Bruceantin | HitBit | 88 | -1a | -100 | CTG |  | 4 | 0 |  | MYC Expression Inhibitor, Apoptosis Inducers |
|  | HLA-ABC | 36 | -1a | -122 | Cell count |  | 4 | 0 |  |  |
|  | HLAB | 370 | -1b | -52 |  |  |  |  |  |  |
| Fluocinonide | HitBit | 4 | -1b | -25 | CTG |  | 4 | 0 | FDA | Glucocorticoid steroid |
|  | HLA-ABC | 13 | -1b | -28 | Cell count |  | 4 | 0 |  |  |
|  | HLAB | 15 | -1b | -18 |  |  |  |  |  |  |
| Methylprednisolone | HitBit | 14 | -1b | -23 | CTG |  | 4 | 0 | FDA | Glucocorticoid steroid |
|  | HLA-ABC | 10 | -1b | -25 | Cell count |  | 4 | 0 |  |  |
|  | HLAB | 24 | -1b | -20 |  |  |  |  |  |  |
| Prednisolone | HitBit | 35 | -1b | -20 | CTG |  | 4 | 0 | FDA | Glucocorticoid steroid |
|  | HLAB | 29 | -1b | -14 |  |  |  |  |  |  |
| Selinexor | HitBit | 630 | -1b | -63 | CTG |  | 4 | 0 | FDA | Exportin-1 Antagonist |
|  | HLA-ABC | 320 | -1a | -100 | Cell count |  | 4 | 0 |  |  |
|  | HLAB | 26,000 | -2b | -60 |  |  |  |  |  |  |
| WS-3 | HitBit | 440 | -1a | -100 | CTG |  | 4 | 0 |  | beta Cell Proliferation Inducer |
|  | HLA-ABC | 65 | -1a | -100 | Cell count | 16,000 | -2b | -100 |  |  |
|  | HLAB | 146 | -1a | -104 |  |  |  |  |  |  |

CC - curve class, IC50 - half max inhibitory concentration

**Supplementary Table 9.** Selective activity for HiBit and HLA-ABC protein

| Sample Name | Activity |  |  |  | Cytotoxicity |  |  |  | Approval | Targets/Mechanism |
| --- | --- | --- | --- | --- | --- | --- | --- | --- | --- | --- |
|  | Data Type | IC50 (nM) | CC | Efficacy | Data Type | IC50 (nM) | CC | Efficacy |  |  |
| ARV-771 | HitBit | 390 | -1a | -100 | CTG |  | 4 | -15 |  | BRD4 inhibitor |
|  | HLA-ABC | 150 | -1a | -100 | Cell count | 29,000 | -3 | -144 |  |  |
|  | HLAB | 160 | -1b | -72 |  |  |  |  |  |  |
| ARV-825 | HitBit | 97 | -1b | -71 | CTG |  | 4 | -20 |  | BRD4 inhibitor |
|  | HLA-ABC | 91 | -1a | -82 | Cell count |  | 4 | 0 |  |  |
|  | HLAB | 65 | -1b | -43 |  |  |  |  |  |  |
| AZD-5153 6-Hydroxy-2-naphthoic acid | HitBit | 130 | -1b | -62 | CTG |  | 4 | 0 |  | BRD4 inhibitor |
|  | HLA-ABC | 14 | -1b | -63 | Cell count |  | 4 | 0 |  |  |
|  | HLAB | 230 | -2b | -69 |  |  |  |  |  |  |
| Bortezomib | HitBit | 31 | -1a | -81 | CTG | 31 | -1b | -37 | FDA | Proteasome Inhibitor |
|  | HLA-ABC | 26 | -1b | -78 | Cell count |  | 4 | 0 |  |  |
|  | HLAB | 820 | -2b | -68 |  |  |  |  |  |  |
| Carfilzomib | HitBit | 88 | -1a | -100 | CTG | 50 | -1b | -38 | FDA | Proteasome Inhibitor |
|  | HLA-ABC | 65 | -1a | -120 | Cell count | 46,000 | -3 | -52 |  |  |
|  | HLAB | 1,800 | -2b | -99 |  |  |  |  |  |  |
| CC-885 | HitBit | 8 | -1a | -100 | CTG |  | 4 | -12 |  | Cereblon (CRBN) Modulator |
|  | HLA-ABC | 8 | -1a | -114 | Cell count |  | 4 | 0 |  |  |
|  | HLAB |  | 4 | 0 |  |  |  |  |  |  |
| CPI-203 | HitBit | 560 | -1b | -58 | CTG |  | 4 | 0 |  | BRD4 inhibitor |
|  | HLA-ABC | 160 | -1b | -60 | Cell count |  | 4 | 0 |  |  |
|  | HLAB | 13 | -1b | -47 |  |  |  |  |  |  |
| Ixazomib citrate | HitBit | 310 | -1b | -74 | CTG |  | 4 | -29 |  | Proteasome Inhibitor |
|  | HLA-ABC | 810 | -1a | -100 | Cell count |  | 4 | 0 |  |  |
|  | HLAB | 6,500 | -2b | -39 |  |  |  |  |  |  |
| OTX-015 | HitBit | 880 | -1b | -63 | CTG |  | 4 | 0 |  | BRD2/4 inhibitor |
|  | HLA-ABC | 230 | -1b | -71 | Cell count |  | 4 | 0 |  |  |
|  | HLAB | 37,000 | -2b | -40 |  |  |  |  |  |  |
| Ouabain | HitBit | 56 | -1a | -83 | CTG | 50 | -1b | -49 | PMDA | Na+/K+-ATPase Inhibitors |
|  | HLA-ABC | 65 | -1a | -81 | Cell count |  | 4 | 0 |  |  |
|  | HLAB |  | 4 | -34 |  |  |  |  |  |  |
| Proscillaridin | HitBit | 12 | -1a | -81 | CTG | 20 | -1b | -51 | PMDA | Na+/K+-ATPase Inhibitors |
|  | HLA-ABC | 36 | -1a | -100 | Cell count |  | 4 | 0 |  |  |
|  | HLAB |  | 4 | 10 |  |  |  |  |  |  |
| SNX-2112 | HitBit | 70 | -1b | -66 | CTG | 560 | -1b | -40 |  | HSP90 Inhibitor |
|  | HLA-ABC | 32 | -1b | -73 | Cell count |  | 4 | 0 |  |  |
|  | HLAB | 23 | -1b | -43 |  |  |  |  |  |  |
| Tanespimycin | HitBit | 70 | -1b | -68 | CTG | 560 | -1b | -40 |  | HSP90 Inhibitor |
|  | HLA-ABC | 13 | -1b | -70 | Cell count |  | 4 | 0 |  |  |
|  | HLAB |  | 4 | 0 |  |  |  |  |  |  |
| TAS-116 | HitBit | 630 | -1b | -63 | CTG | 7,000 | -1b | -51 |  | HSP90 Inhibitor |
|  | HLA-ABC | 260 | -1a | -83 | Cell count |  | 4 | 0 |  |  |
|  | HLAB | 370 | -1b | -30 |  |  |  |  |  |  |
| Thapsigargin | HitBit | 8 | -1a | -100 | CTG | 4 | -1b | -37 |  | SERCA inhibitor |
|  | HLA-ABC | 81 | -1b | -76 | Cell count | 14 | -3 | -22 |  |  |
|  | HLAB | 29,000 | -3 | -128 |  |  |  |  |  |  |

CC - curve class, IC50 - half max inhibitory concentration

**Supplementary Table 10.**
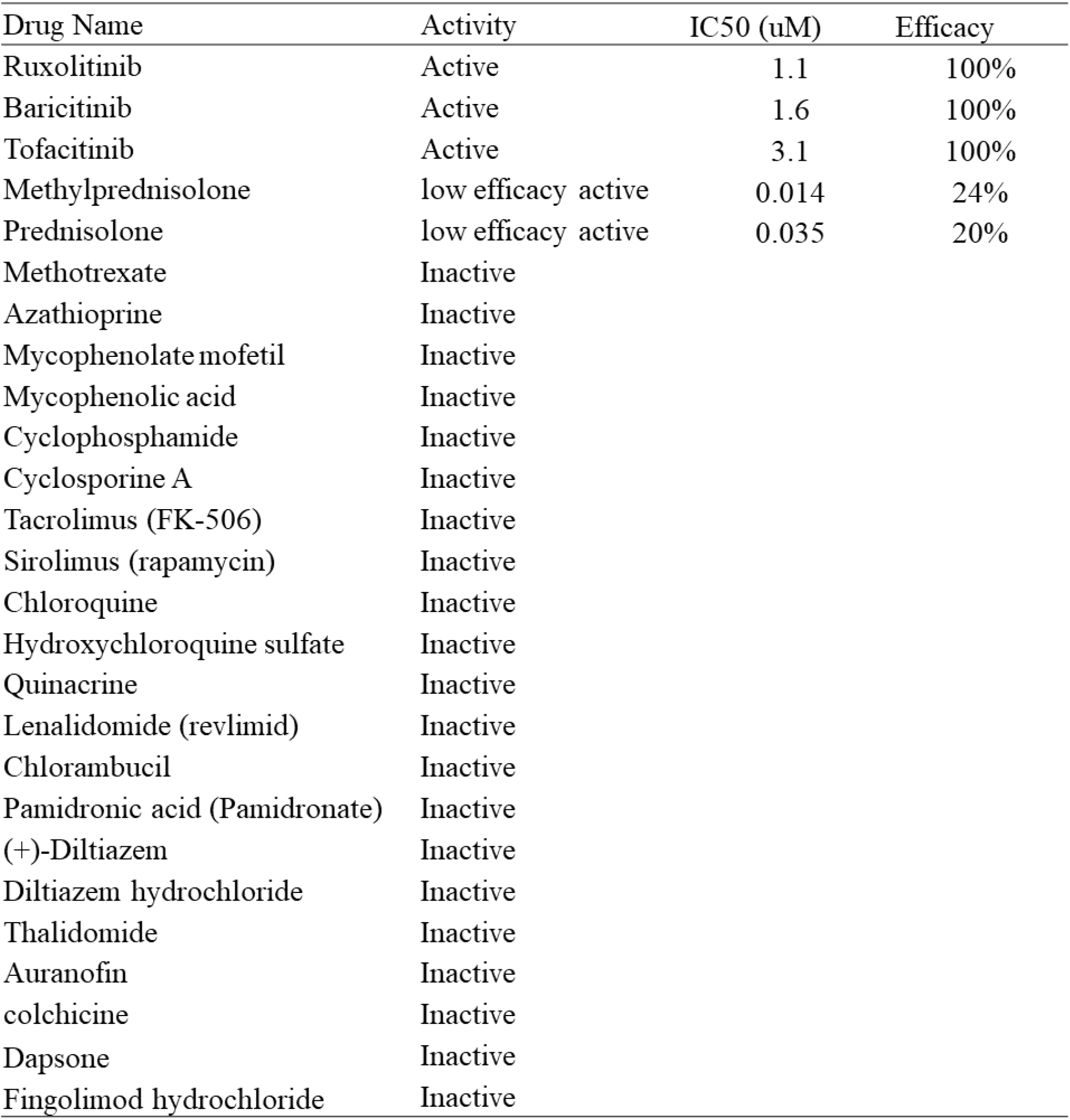
Primary HLA-B Hi.Bit screenin g activities of small molecule myositis therapeutics

## SUPPLEMENTARY MATERIALS AND METHODS

### HLA-ABC IF secondary assay

All aspiration and dispensing steps for immunostaining were conducted at ambient temperature with an EL406 (BioTek, Winooski, VT) with a 16×4 pin aspiration head and an 8 tip, 5 uL dispensing cassette. All solutions were prepared in PBS. Cells were fixed in 4.0% paraformaldehyde for 20 min, permeabilized in 0.1% Triton X-100 for 20 min, blocked in 10% goat serum for 30 min, incubated with 1:400 HLA-ABC antibody (clone W6/32, MA5-11723) + 2% goat serum + .05% tween-20 for 2.5 hrs, washed three times, incubated with 1:1000 Hoechst 33342 (H3570) + 1:1000 Alexa Fluor 488 goat anti-mouse antibody (A11001) + 1:5000 CellMask Red (H32712) for 1 hr, washed three times, and stored at 4 °C for later analysis

### Western blotting

All lysates were stored at −80 °C for later analysis. Thawed samples were cleared of debris by centrifugation, and supernatant protein concentration was determined by BCA assay (23227).

For HiBit clones, 10 ug of protein/sample were prepared and reduced in NuPAGE LDS Sample buffer (NP00008) + 5% BME, then run on 1.0 mm, 12-well, 4-12% NuPAGE Bis-Tris gels in MOPS running buffer (NP0322 and NP0001) at 150 V for 80 min, and transferred to 0.45 um pore PVDF membrane (Bio-Rad, Hercules, CA, Immun-Blot LF, 1620263) at 150 mA for 1 hr in NuPAGE Transfer Buffer (NP0006) with 10% methanol. Lane 1 contained Amersham ECL Plex Fluorescent Rainbow Marker, Full Range (RPN850E). All blocking and immunoblotting was done with 5% non-fat milk powder in tris buffered saline + 0.05% tween-20 (TBST). Membranes were incubated with a 1:2000 mouse HLA-ABC antibody (HC-10, gift from Dr. Robert A. Colbert at National Institute of Arthritis and Musculoskeletal and Skin Diseases) + 1:1000 rabbit β-tubulin antibody (abcam, ab179513) overnight at 4 °C. After three washes in TBST, membranes were incubated with Alexa Fluor 647 goat anti-mouse antibody (A21235) for HLA-ABC and Alexa Fluor 488 goat anti-rabbit antibody (A1108) for β -tubulin, both diluted 1:2000 for 1 hr at ambient temperature. After three washes in TBST, membranes were imaged on a Typoon FLA 9500 (GE). Membranes were incubated with Restore Stripping Buffer (21059) for 15 min, washed 3 times in TBST, then HiBit was detected with the Nano-Glo HiBit Blotting System (Promega, N2410) according to the manufacturer’s protocol using a 1 hr incubation with LgBit and chemiluminescent detection on a Bio-Rad Chemidoc Touch.

For active compound 30 min treatment, parental C25cl48 cells were pretreated with compounds for 1 hr prior to adding IFN-β for 30 min, whereas for the 24 hr treatment, compounds and IFN-β were added at same time. For both experiments, compounds were tested at their IC95 according to the HiBit CRCs (in uM): Rux, Bari, and Tofa – 6.3, Pred – 1.0, Echi - 0.032, Givi – 18.2, Pano - 0.079, CUD – 1.0, Dina - 0.079, ARV – 0.13, Nemo – 0.32. For Western blot, 15 ug of protein/sample were prepared and reduced in NuPAGE LDS Sample buffer (NP00008) + 5% BME, then run on 4-12% Bis-Tris gel, 10×8, 15-well (GenScript, Piscataway, NJ, M00654) at 150 V for 60 min, and transferred to 0.45 um pore nitrocellulose (Bio-Rad) at 150 mA for 1 hr in NuPAGE Transfer Buffer (NP0006) with 10% methanol. Lane 1 contained Amersham ECL Plex Fluorescent Rainbow Marker, Full Range. Membranes were blocked in 5% bovine serum albumin (BSA) in TBST, and antibodies were prepared in 1% BSA in TBST and incubated for 1 hr at ambient temperature except phospho-STAT1 was overnight at 4 °C. Antibodies used were 1:1000 rabbit phospho-STAT1 antibody (Cell Signaling Technology, Danvers, MA, 7649), 1:1000 rabbit STAT1 antibody (Cell Signaling Technology, 9172T), 1:1000 mouse HLA-ABC antibody (HC-10), and 1:1000 rabbit β-tubulin antibody. After three washes in TBST, membranes were incubated with either 1:5000 goat anti-rabbit stabilized peroxidase secondary antibody (Invitrogen 32460) or 1:5000 goat anti-mouse stabilized peroxidase secondary antibody (Invitrogen 32430) for 1 hr at ambient temperature. After three washes in TBST, membranes were imaged using Super Signal West Dura (34076) and a BioRad Chemidoc Touch. Membranes were stripped between each primary + secondary antibody pair as described above.

### Whole Genome Sequencing

For WGS library preparation, an average of 200 ng of quality DNA was used. The sequencing libraries were made using the Twist library-enzymatic protocol according to manufacturer’s instructions (Twist Biosciences, CA, USA). The fragmentation conditions were adapted to create fragments around 300-400bp followed by dual-SPRI size selection washing procedure. In summary gDNA was utilized for Fragmentation, End Repair, dA-tailing and Adapter Ligation. Twist Universal Adapter system consists of Twist Universal Adapters and Twist Unique Dual Indexed (UDI) Primers which provides high quality libraries. Two PCR cycles were applied after library addition to increase the yield and close the molecule ends for better quality control check on a Bionalyzer and qPCR. The duplicate rates were minimal and did not really impact the data with this library method. WES sequencing for 5 samples was performed on the Novaseq6000 (Illumina, La Jolla, CA, USA) using the S4 Flow cell with the paired-end 300 cycle protocol (2×150bp) following Illumina recommendations.

Raw sequencing data was outputted by the Illumina NovaSeq 6000 in the form of BCL (binary base call) files. The Illumina DRAGEN Bio-IT Platform (version 3.3.7) was used to generate a final output of VCF (variant call format) files for each sample. In more detail, the BCL files are first demultiplexed and converted into FASTQ files. DRAGEN then aligns these FASTQ files to the Homo sapiens GRCh37 genome (hs37d5) to generate CRAM alignment files. During this step, the alignment file was also sorted and duplicate reads marked. The final step performed by DRAGEN was the variant caller which generated a VCF file from the CRAM input and contained single-nucleotide variants (SNVs) and short insertions/deletions (indels). The HiBit insertion sites in the test samples were identified by comparing the short insertions in the VCF files from the control and test samples. The control cells have two heterozygous alleles for HLA-B (08:01 and 18:01). HiBit sequence was designed to target the HLA-B 08:01 allele. The HLA-B 08:01 allele contains a single-nucleotide variant (dbSNP:rs2308655 C>G) which is not present in HLA-B 18:01 and is 40bp away from the designed insertion site. The phase of the rs2308655 variant and the HiBit insertion site can be determined by checking whether they exist on the same short read (read-backed phasing).

### Protocol Tables for qHTS

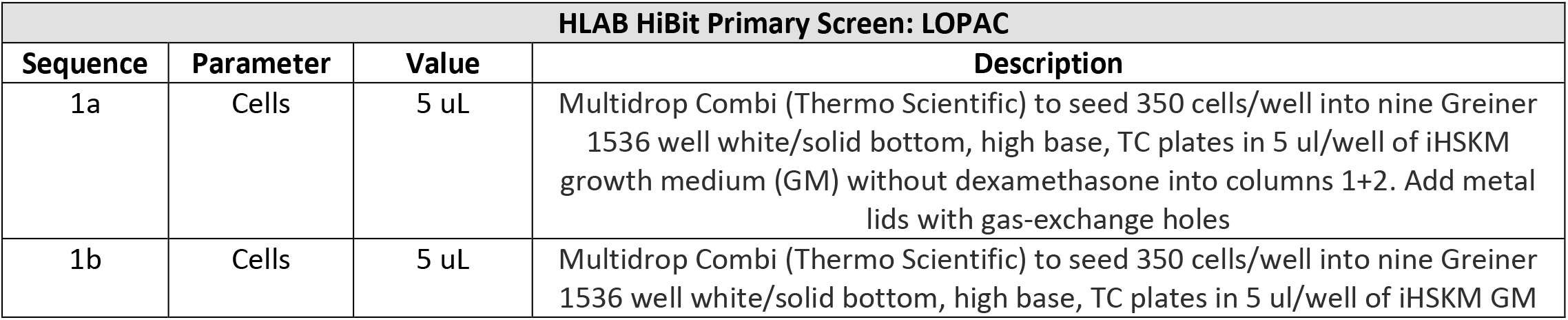
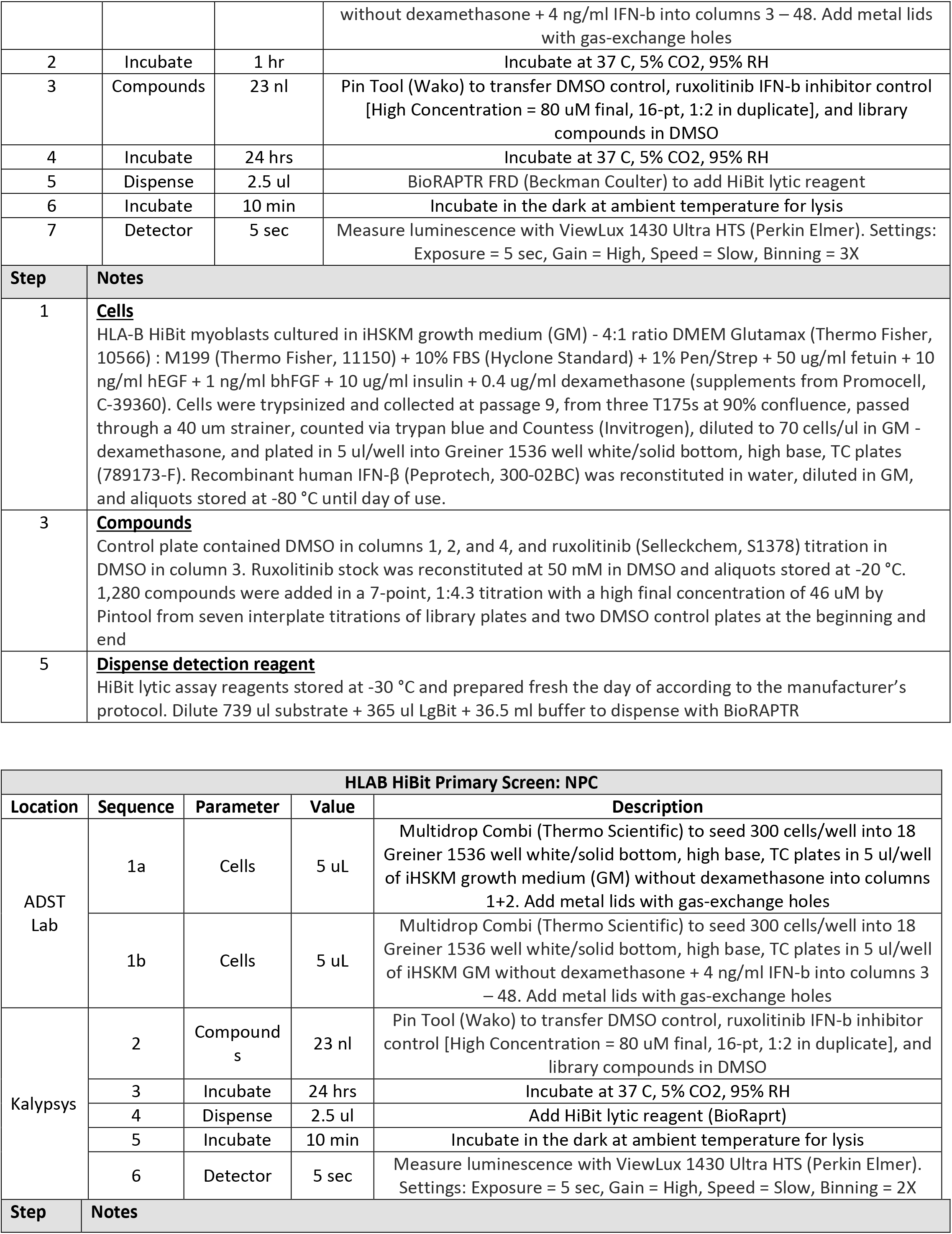
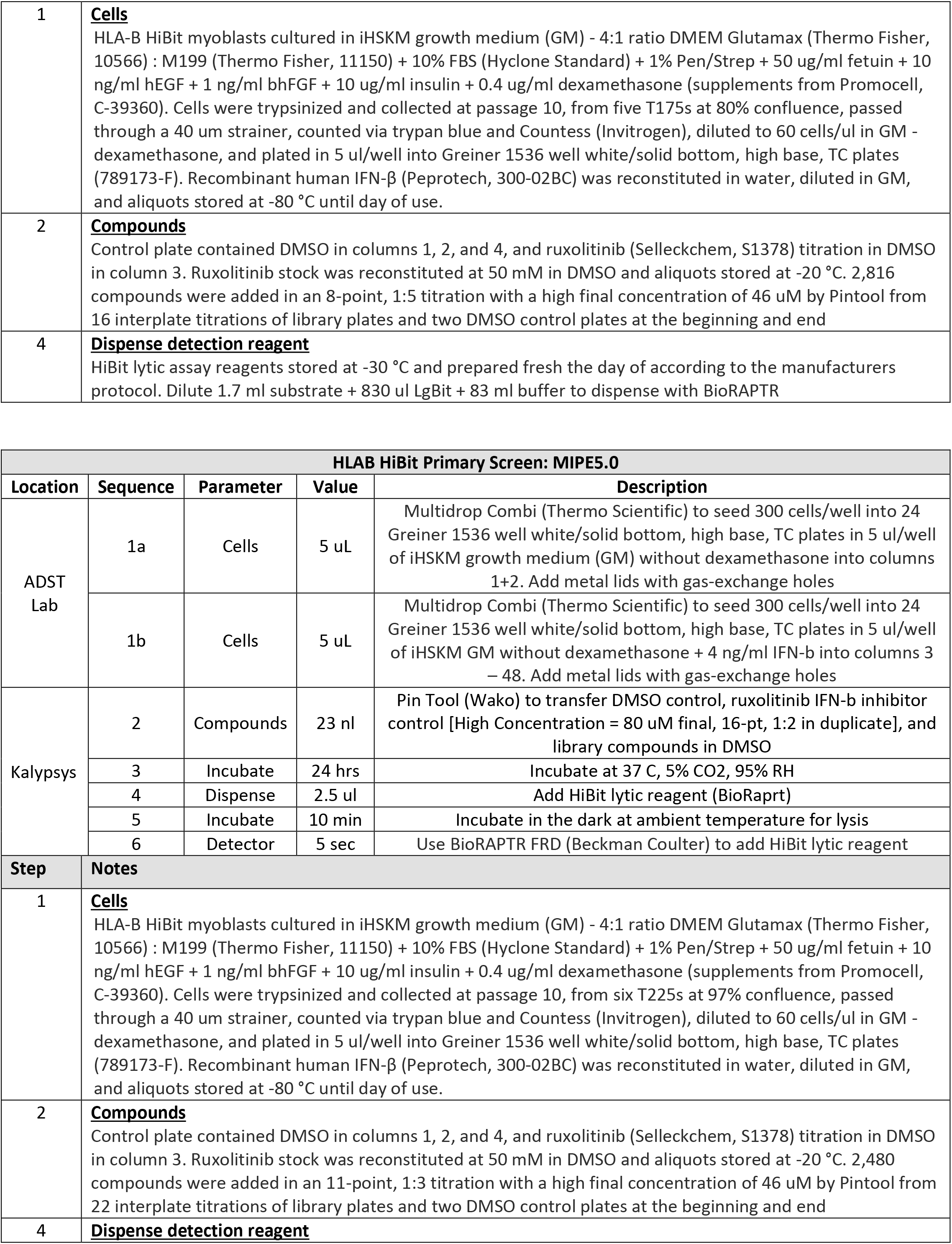
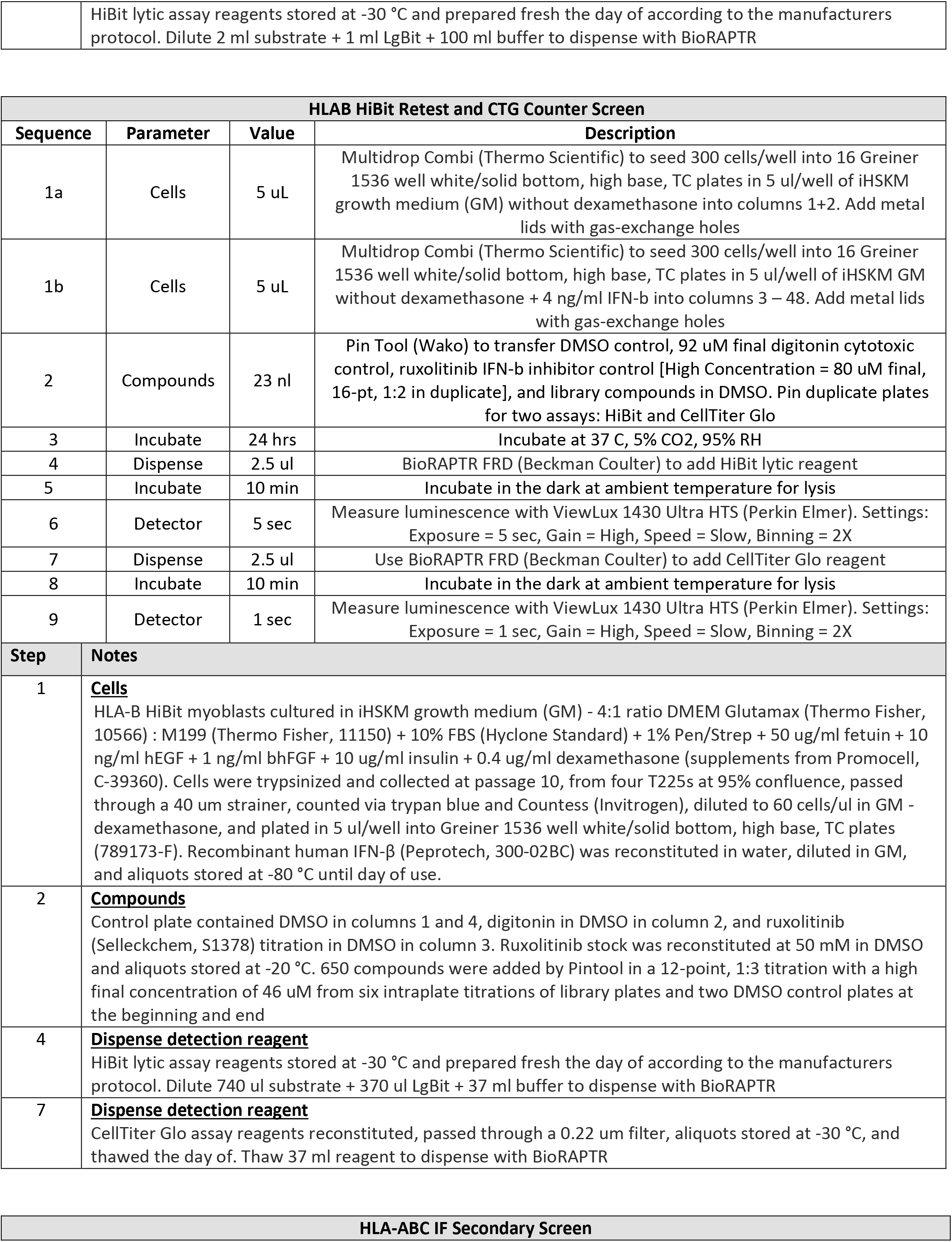
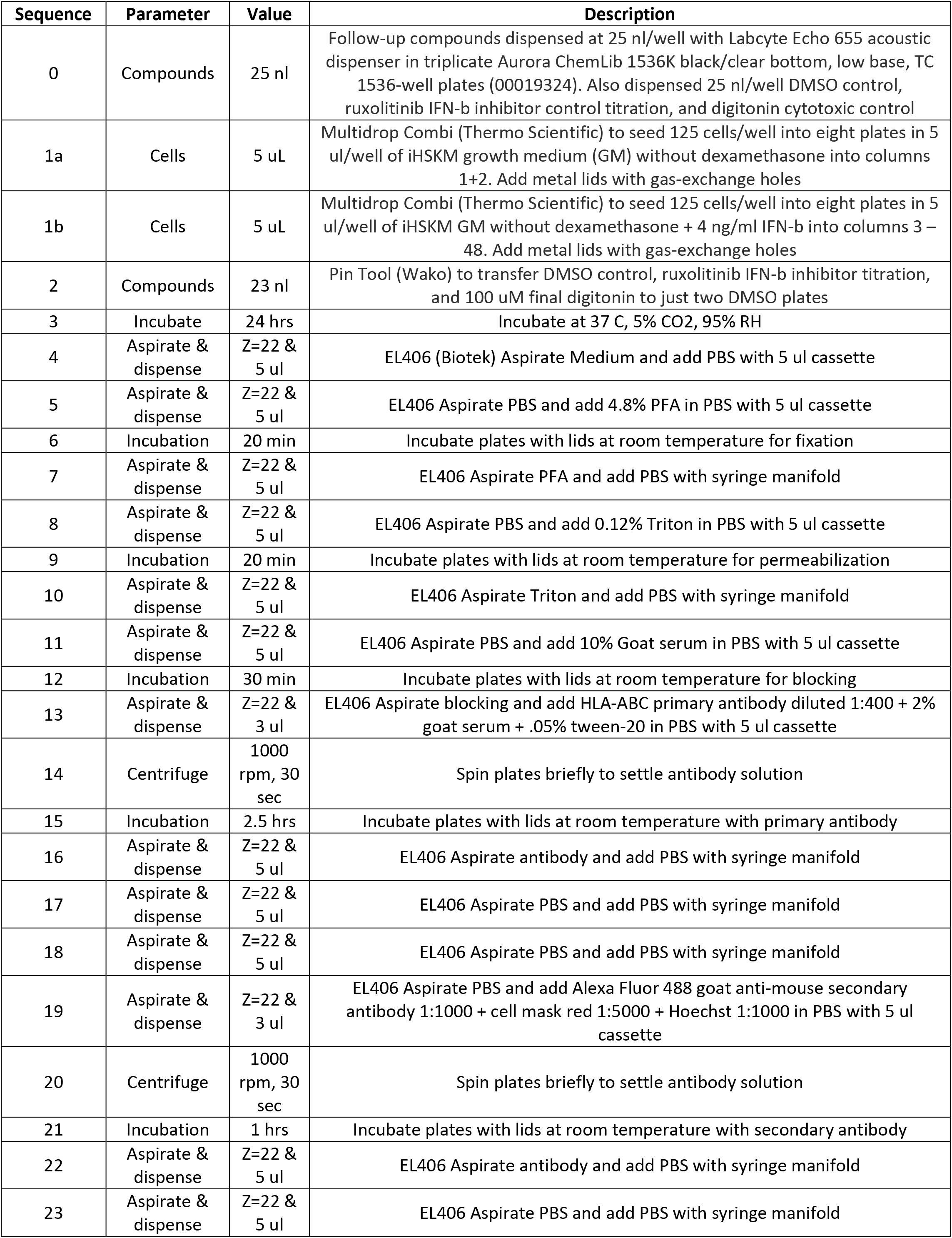
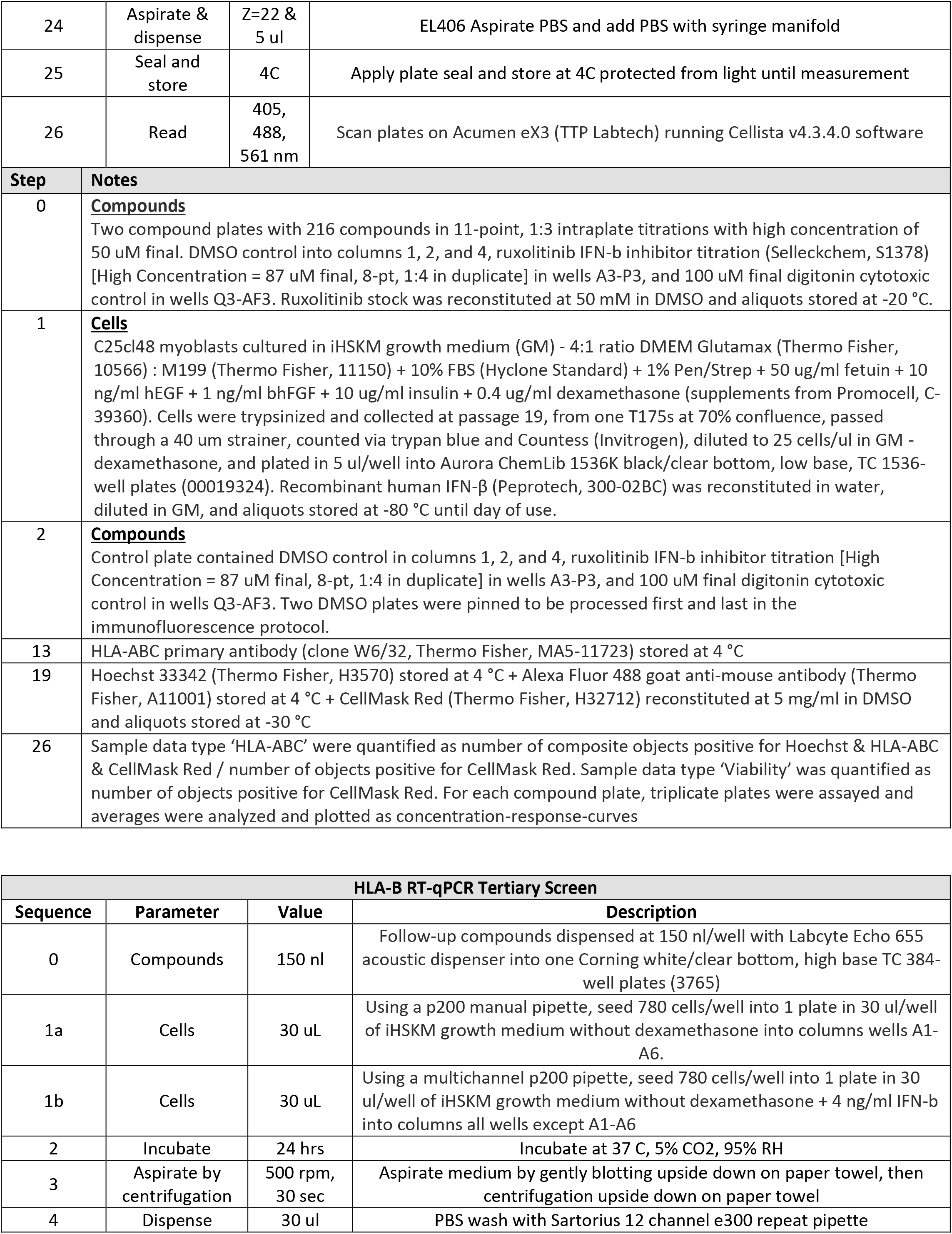
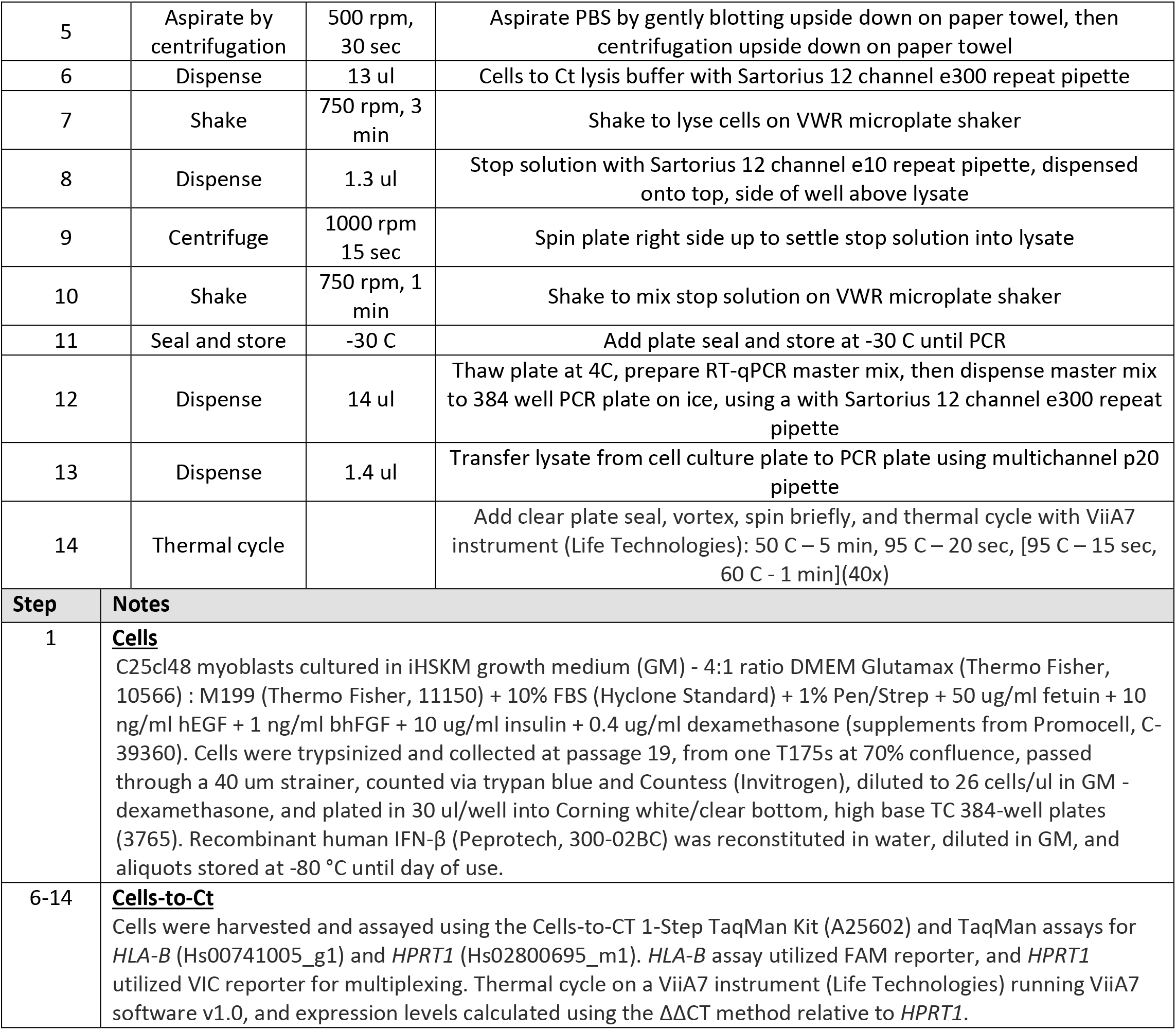

